# Isolation of cardiomyocytes undergoing mitosis with complete cytokinesis

**DOI:** 10.1101/549212

**Authors:** Hsiao-yun Y. Milliron, Matthew J. Weiland, Eric J. Kort, Stefan Jovinge

**Author notes:** Equal contribution (Hsiao-yun Y. Milliron, Matthew J. Weiland, Eric J. Kort). H.Y.M. and S.J. designed the study. H.Y.M. and M.J.W. performed the experiments. H.Y.M., M.J.W. and E.J.K. performed analysis. H.Y.M., M.J.W., E.J.K. and S.J. wrote the manuscript.

## Abstract

Rationale-Adult human cardiomyocytes (CMs) do not complete cytokinesis despite passing through the S-phase of the cell cycle. As a result polyploidization and multinucleation occur. In order to get a deeper understanding of the mechanisms surrounding division of CMs there is a crucial need for a technique to isolate CMs that complete cell division/cytokinesis.

Objective-Markers of cell cycle progression based on DNA content cannot distinguish between mitotic CMs that fail to complete cytokinesis from those cells that undergo true cell division. With the use of molecular beacons (MB) targeting specific mRNAs we aimed to identify truly proliferative CMs derived from hiPSCs.

Methods and Results-Fluorescence activated cell-sorting combined with molecular beacons was performed to sort CM populations enriched for mitotic cells. Expressions of cell-cycle specific genes were confirmed by means of RT-qPCR, single-cell RNA sequencing (scRNA-seq). We further characterized the sorted groups by proliferation assays and time-lapse microscopy which confirmed the proliferative advantage of MB-positive cell populations relative to MB-negative and G2/M populations. Gene expression analysis revealed that the MB-positive CM subpopulation exhibited patterns consistent with the biological processes of nuclear division, chromosome segregation, and transition from M to G1 phase. The use of dual-MBs targeting *CDC20* and *SPG20* mRNAs (*CDC20*^+^*SPG20*^+^) enabled the enrichment of cytokinetic events. Interestingly, cells that did not complete cytokinesis and remained binucleated were found to be *CDC20*^−^*SPG20*^+^ while polyploid CMs that replicated DNA but failed to complete karyokinesis were found to be *CDC20*^−^*SPG20*^−^.

Conclusions-This study demonstrates a novel alternative to existing DNA content-based approaches for sorting CMs with true mitotic potential that can be used to study in detail the unique dynamics of CM nuclei during mitosis. Together with high-throughput scRNA-seq, our technique for sorting live CMs undergoing cytokinesis would provide a basis for future studies to uncover mechanisms underlying the development and regeneration of heart tissue.

## Introduction

Myocardial infarction (MI) remains one of the leading causes of death in the United States and in most industrialized nations throughout the world.^1^ In-hospital mortality rates of patients suffering an MI have improved as a result of major advances in cardiovascular science and medicine. However, survivors of MI have an increased risk of developing heart failure and have high rates of early post-discharge death and re-hospitalization.^3–5^ While mortality from coronary artery disease has declined, heart failure has emerged as a leading public health burden worldwide.^4,6^ Ischemia associated cardiomyopathy following a myocardial event is characterized by CM loss and failure of myocardial regeneration. While we have demonstrated that the post-natal human heart retains a modest proliferative potential, ^7,8^ the endogenous proliferative capacity of the adult heart remains insufficient for recovery of this CM loss.

It is well known that a majority of postnatal human CMs undergo repeated rounds of DNA replication without cytokinesis resulting in polyploidy and multinucleation.^7,9–11^ As the human heart matures, the endogenous proliferative capacity diminishes and nonproductive cell cycle activities lead to the formation of binucleated and polyploid daughter cells. The percentage of diploid CMs decreases steadily with a concomitant rise in polyploid cells, with some cells reaching 8N or higher. While this underscores the fact that CMs have substantial capacity for DNA replication,^9,12^ a unique experimental challenge exists with respect to identifying CMs that complete cytokinesis. Due to their propensity for polyploidization and multinucleation, markers of cell cycle progression based on DNA content such as Ki67 or thymidine analogues cannot distinguish between merely polyploid CMs and those cells that are in the process of true cellular division. Moreover, assays based solely on DNA content do not reveal information on cell stage within M phase of the cell cycle (Fig. 1a,b). Since DNA content alone does not identify CMs expressing late mitotic markers, novel experimental strategies for identification, sorting, and enrichment of these proliferating cells are required.

**Figure 1.**
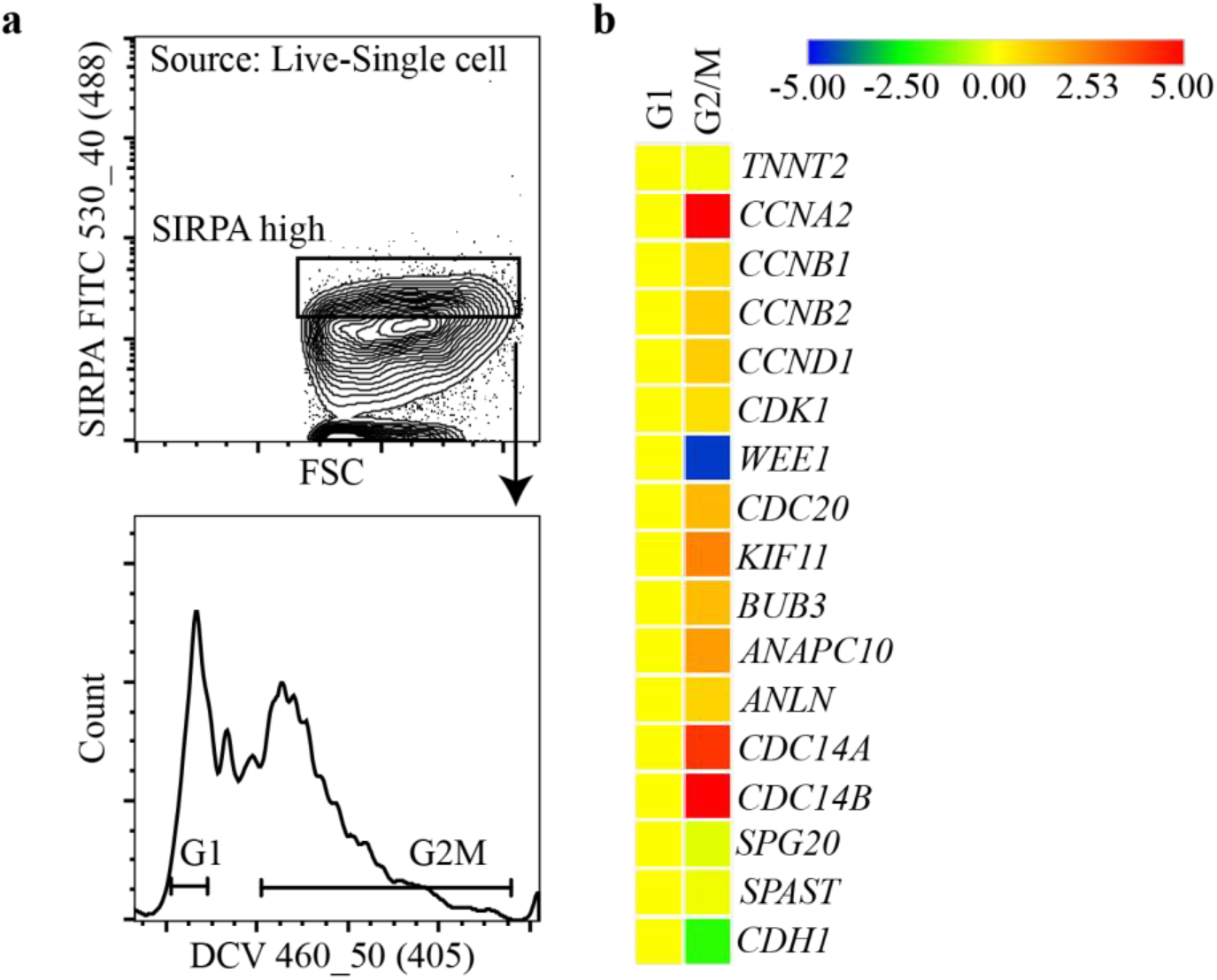
FC sorting of D8 hiPSC-CMs by DNA content does not significantly enrich for CMs in late M phase. **a)**Representative FC plots of D8 hiPSC-CMs stained with SIRPA and DNA content dye DyeCycle Vybrant Violet and the cell sorting strategy of FSC/SSC, singlets, viable, SIRPA^high^ and G1 and G2/M. b)RT-qPCR of 50 sorted events from the G1 and G2/M gated populations. Heatmap showing the Z-score of ΔΔCt values of G2/M normalized to G1 (set to 1). The heatmap was generated using Morpheus.

Based on quantification of ^14^C incorporation into myocardial DNA, we have previously demonstrated that the annual CM turnover rate decreases from 1-2% per year at the age of 25 to 0.45% per year at the age of 75.^7,8^ Due to this turnover, approximately 40% of CMs in individuals at 70 years of age have been generated post-birth.^7^ Furthermore, we recently reported that these new CMs are derived from pre-existing cytokinetic CMs.^13,14^ A promising strategy to achieve recovery after MI would be enhancement of this endogenous proliferative potential retained by the myocardium. To better understand how to unlock this process, we must first have methods to identify and isolate live proliferating CMs.

Molecular beacons (MBs) allow for RNA detection in live cells. MBs are single-stranded oligonucleotides possessing a stem-loop structure. The loop contains a probe sequence that is complementary to a target sequence, while the stem structure is composed of complementary base pairs that bring a fluorophore and quencher into close proximity. In the presence of a complementary target sequence, the beacon undergoes a conformational change restoring fluorescence.^15–17^ It has been demonstrated that MBs can be used in flow cytometry (FC) assays to identify cells expressing specific mRNA transcripts within a mixed cell population without disrupting gene expression^18^ and this approach has been used previously to identify CM subtypes in other contexts.^19–22^ As this strategy allows sorting of live cells, cells may then be used for downstream studies or clinical applications. In this study, we employ human iPSC-derived CMs which are the best current *in vitro* model of CM cell biology. While these cells in many ways resemble fetal-like CMs, we also tested our hypotheses in cells aged out to increasing levels of maturity, and cross reference our findings with a rat-CM binucleation model.

The cell-division cycle protein 20 (CDC20) is required to activate the anaphase promoting complex initiating chromatid separation and entrance into anaphase, leading to Securin and Cyclin B degradation and eventually mitotic exit.^23–26^ Therefore we hypothesized that the level of *CDC20* mRNA in live cells would be a candidate marker to indicate cell cycle status of cells in late mitosis. Likewise, the Microtubule Interacting and Trafficking molecule domain-associated gene, *SPG20*, is associated with the Endosomal Sorting Complexes Required for Transport (ESCRT) machinery and cytokinesis. The *SPG20* gene encodes the protein spartin which is involved in the ESCRT pathway and midbody abscission during completion of cell division.^27–29^ Here we investigate to what extent these markers-individually or together-can be targeted with MB technology to enrich cells undergoing anaphase and telophase.

## Methods

### An expanded Methods section is available in the Online Data Supplement

### Differentiation and long-term maintenance of hiPSC-CMs

Episomal human iPSCs^30,31^ (Thermo Fisher Scientific) were maintained as a monolayer on Matrigel (Corning Life Sciences) coated plates and supplemented with RPMI 1640 (Gibco) containing B27 minus insulin (RPMI/B27-insulin) with GSK3 inhibitor CHIR990216 at D0-1 to induce mesoderm, followed with cardiac differentiation at D4-5 by changing media to RPMI 1640/B27 (-insulin) with Wnt inhibitor IWR1.^31^ For long-term hiPSC-CM maintenance, cells were treated with RPMI 1640 (-L-glutamine and -glucose) (Biological Industries USA, Inc.) containing B27 with insulin (RPMI/B27 +insulin) at D10-13. At D13, cells were returned to RPMI/B27 +insulin and stabilized for 48h. Cells were then dissociated and replated onto a new Matrigel-coated well. Media was replaced with fresh RPMI/B27 +insulin every three days.

### Isolation of target cells using flow based on MB signal

For MBs complementary to human-specific *CDC20* or *SPG20* mRNAs, multiple candidate target sequences were designed for each target gene and tested for fluorescence shift prior to cell sorting for cellular amplification of mRNA targets. Probes with the highest fluorescence were considered to pass the flow QC (quality control) and subjected to the RT-qPCR checkpoint. In order to facilitate cellular delivery, all MBs were modified with the cell-penetrating peptide MPG (GALFLGFLGAAGSTMGAWSQPKSKRKV, with N-terminal acetylation and C-terminal cysteamide modification) to form a stable non-covalent complex and promote the uniform and rapid delivery of MBs into cells without interfering with specific targets or hybridization-induced fluorescence.^32,33^ Cells were then added to the MB/MPG complex medium for MB-target hybridization and analyzed/sorted on a BD Influx (BD Biosciences) flow cytometer.

### Generation of single cell GEMs and sequencing libraries

After sorting, downstream cDNA synthesis and library preparation were carried out as instructed by the manufacturer^34^ (Chromium Single Cell 3’ Library & Gel Bead Kit v2). Libraries were standardized for cDNA mass, pooled, and sequenced on an Illumina NextSeq 500 sequencer. Base calling was done by Illumina NextSeq Control Software (NCS) v2.0 and output of NCS was demultiplexed and converted to FastQ format the Cell Ranger mkfastq pipeline (10X Genomics).

### Bioinformatics analysis of scRNA-seq

Demultiplexed samples were aligned to the reference human genome (version GRCh38) and converted to transcript counts using the Cell Ranger pipeline. The AUCell algorithm^35^ was used to perform gene set enrichment analysis. Differences between groups with respect to the AUCell scores were evaluated by t-test. We used the pagoda2 package^36^ to identify genes that were differentially expressed between cell populations. Enrichment for gene ontology terms was calculated with the DOSE package.^37^

## Results

### MB-MPG delivery and selection of optimal MBs for isolating hiPSC-CMs at M-phase

To differentiate cells in anaphase/telophase from those in earlier stages of mitosis, MBs targeting *CDC20* mRNAs were designed to identify mitotic events where the onset of anaphase was likely to occur. *CDC20* acts as a key activator promoting metaphase to anaphase transition by converging on the SAC,^26,38,39^ and we hypothesized that truly proliferative (as opposed to polyploid and binucleated) CMs could be identified based on expression of mRNA encoding markers of late mitosis (*CDC20* and *SPG20*). To accomplish this, we first optimized MB-MPG delivery by evaluating the cell viability, MB background due to endogenous nuclease degradation, and the delivery efficiency by using a positive control MB lacking its quencher (Fig. 2b**; Online Table 1**). The MPG sequence used in this study has a variant characterized by a single mutation of the second residue in the nuclear localization sequences (NLS) motif, from lysine to serine (KSKRKV instead of KKKRKV). This mutation has previously been reported to dramatically reduce the import of NLS-containing proteins into the nucleus.^33,40,41^ As shown in Fig. 2d, the transduction efficiency of ~90% was achieved after 1h incubation with MB-MPG, while maintaining >90% viability. The closely packed contour suggested that the signal shift was due to the cellular uptake of fluorescently labeled MBs (Fig. 2b). Within the established timeframe, we demonstrated that the MB-MPG delivery was completed without appreciable signal resulting from beacon degradation (Fig 2b,d). The intracellular uptake efficiency was also tested by incubating MB-MPG with a monolayer of hiPSC-CMs. Fluorescence was detectable in 100% of the live cells 1h after delivery. Homogeneous distribution of fluorescence in both cytoplasm and nucleus was observed, indicating that MPG is a reliable tool for the delivery of mRNA-targeting MBs (Fig. 2c). We then transduced heterogeneous populations of hiPSC-CMs with MB candidate sequences targeting *CDC20* or *SPG20* mRNAs to further verify the specificity. The cells were separated by FC based on signal shifts (Fig. 3a, 6b). Interestingly, G2/M phase cells clearly showed a higher MB signal shift than cells in G1 phase, which was a further indication that the expression of the target sequences was putatively more abundant in the later phase of cell cycle. Two *CDC20* MBs passed the flow QC as both MBs resulted in a substantially high percentage of fluorescent cells, and the sorted cells were further characterized by RT-qPCR to check the MB specificity (Fig. 3a,b; Online Fig. 1b). In addition to the upregulation of genes involved in cell cycle progression and mitosis, the RT-qPCR analyses also revealed high levels of midbody-associated *SPG20* and *SPAST* (spastin) mRNA expression in sorted *CDC20*^+^ events compared with *CDC20*^−^ events (Fig. 3a,b). The same QC strategies were applied to *SPG20* MBs (Fig. 6b; Online Fig. 1a,b). Based on qRT-PCR results, *CDC20-642* and *SPG20-2264* MBs were chosen for further experiments described in this report.

**Figure 2.**
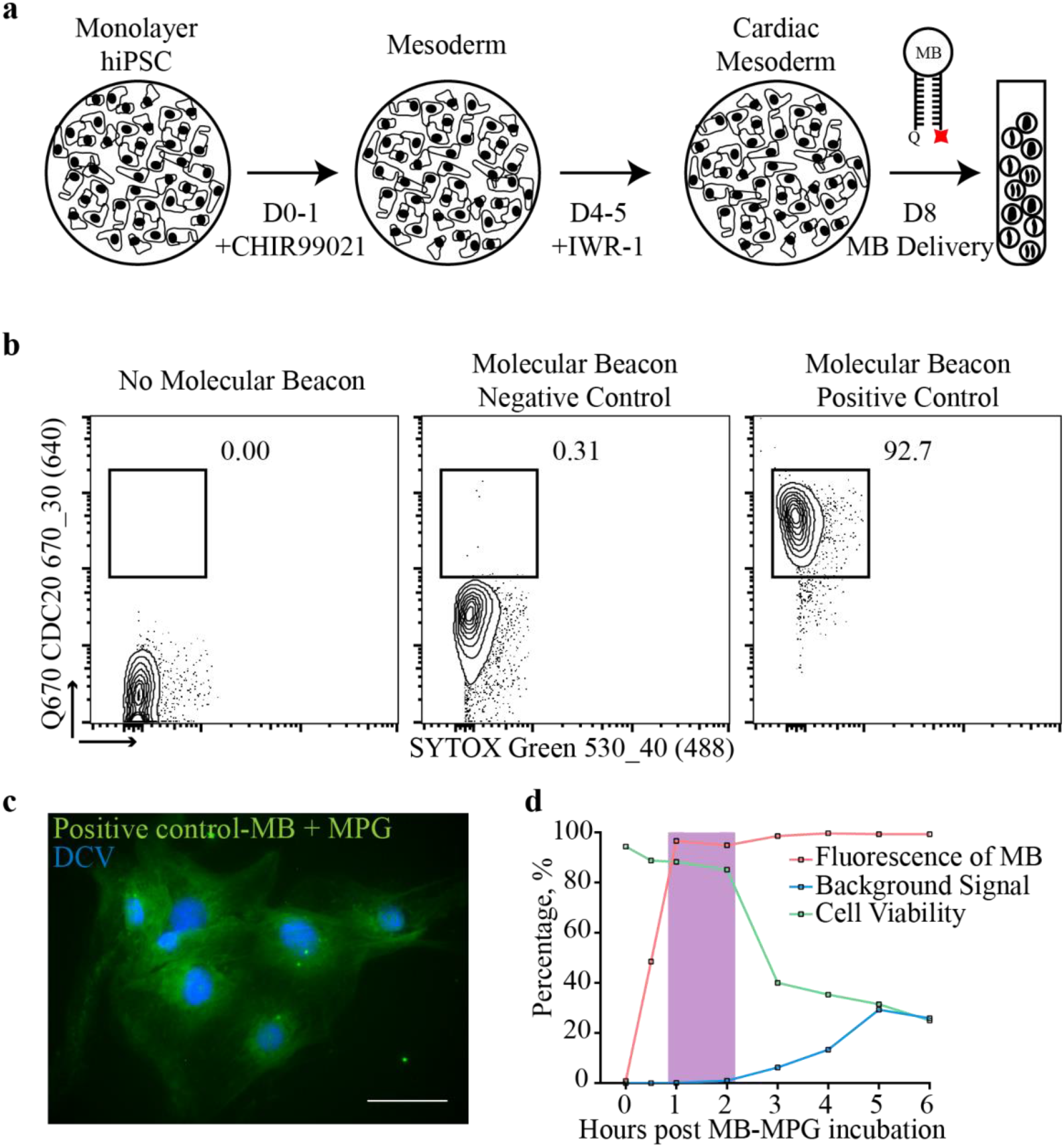
Human iPSC cardiac differentiation and efficient delivery of MPG coupled molecular beacon (MB). **a)**Small molecule hiPSC cardiac differentiation protocol. Cells were harvested at D8 followed by delivery of MBs targeting mRNA of interest and sorted on a flow cytometer. b)Representative FC analysis of the delivery efficiency of MBs. Left panel, live D8 hiPSC-CMs with no MB delivered. Center panel, delivery of a negative control MB. Minimal background (0.31%) was observed after MB-MPG delivery. Right panel, delivery of positive control MB, ensuring fluorescence of MB upon delivery (>90% of D8 hiPSC-CMs). Gating hierarchy: FSC/SSC to trigger pulse width/FSC to SSC-A/SSC to live (SYTOX Green negative) events. Numbers represent percentages of MB positive events. c)Image of delivery of MB-MPG into a monolayer of live D8 hiPSC-CMs. Green-6 FAM-labeled MB-MPG, Blue-DyeCycle Vybrant Violet (DCV). Scale bar, 50μm. d)MB delivery kinetics and viability in D8 hiPSC-CMs. Quantification of delivery efficiency (positive control MB (orange)), background fluorescence (negative control MB (blue)) and cell viability (green) in D8 hiPSC-CMs. Purple indicates optimal delivery time point with the highest fluorescence of positive control MB and the highest cell viability. Background fluorescence of negative control MB was negligible following incubation times of 1 and 2h.

**Figure 3.**
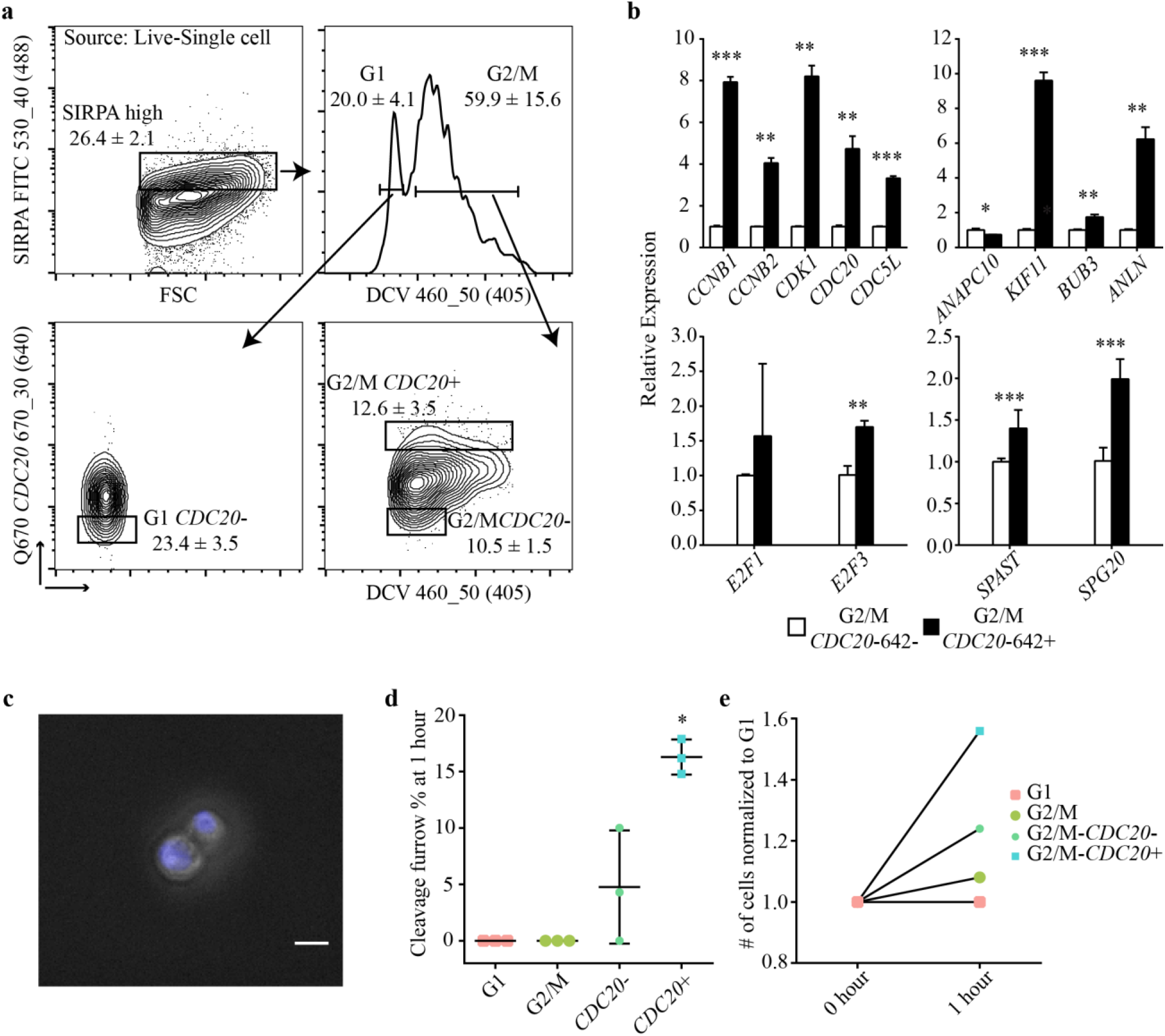
FC sorting of G2/M-*CDC20*^+^ events enrich for D8 hiPSC-CMs in M phase. **a)**Representative FC analyses of the cell sorting strategy used for the isolation of G1 and G2/M, G2/M-*CDC20*^−^ and G2/M-*CDC20*^+^ events following delivery of *CDC20* MB. Gating strategy; Debris, doublets, and non-viable events were excluded, SIRPA^high^, DCV DNA content for G1 and G2/M, *CDC20*^+^ and *CDC20*^−^ events. Values represent mean parental gate percentages ± SD (n = 3). **b)**RT-qPCR analysis of the expression levels of cell cycle, cell cycle regulator, mitosis, proliferation and late M-phase genes from 50 G2/M-*CDC20*^−^ (open bar) and G2/M-*CDC20*^+^ (black bar) sorted events. (t-test: Mean±SD; n = 3; *p < 0.05, **p < 0.01 ***p < 0.001) **c)**Representative images of BF/DCV overlay from G2/M-*CDC20*^+^ events 1h post-sort with a distinct cleavage furrow. Scale bars, 50μm. **d)**Quantifìcation of the percentage of cells with a cleavage furrow at 1h post-sort from the sorted populations. (t-test: Mean±SD; n = 3; *p < 0.05) **e)**Cell count data 1 h post-sort from the sorted populations, values were normalized to G1 (set at 1) (t-test; Mean±SD; n = 3; *p < 0.05)

### Accounting for non-biologic loss of CTY fluorescence

To quantitate the proliferation advantage of G2/M-*CDC20*^+^ CMs relative to G2/M-*CDC20*^−^ and G1-*CDC20*^−^ CMs, we analyzed sorted populations at different post-sort time points to obtain a time course of CTY fluorescence. We found that more than 30% of cells sorted from G1 had a decrease in fluorescence between 0 and 1h (data not shown), which is unlikely a result of cell cycle progression. In addition, we found no increase in cell count from the G1 sorted population. To quantify the passive efflux of CTY, cells were arrested at G2/M with Nocodazole. In the presence of Nocodazole, cells were plated and harvested at 1h and 2h in order to observe fluorescence. Arrested cells had large decreases in fluorescence at both 1 and 2h (**Online** Fig. 2b). Therefore, loss of fluorescence was attributed to a cell specific mechanism other than halving at each division. Since we were confident that G1 cells were not dividing within 18 hours post-sort, we calculated percent dividing cells from G2/M-*CDC20*^+^ and G2/M-*CDC20*^−^ by subtracting G1 percentage at the same time points.

### G2/M-*CDC20*^+^ hiPSC-CMs have a proliferative advantage

To further evaluate whether G2/M-*CDC20*^+^ CMs were enriched for cells in M phase, we isolated them from D8 cultures and compared gene signatures, cellular morphology, cell counts, and proliferation to the G1, G2/M, and G2/M-*CDC20*^−^ cell populations. RT-qPCR analyses of 50 cells showed that G2/M-*CDC20*^+^ populations have a selected gene set enriched (p < 0.05, t-test) for cells actively progressing through the cell cycle and are in the late stages of M phase, including *CDC20* and *SPG20* expression (Fig. 3b).

In animal cells, the cleavage furrow is formed in late mitosis with the appearance of a shallow groove on the surface of dividing cells^42^. Cultures of G2/M-*CDC20*^+^ cells, 1h post-sort, had a significantly higher percentage of cells (16.3±1.55) with a prominent cleavage furrow than G1 (0), G2/M (0), and G2/M-*CDC20*^−^ (4.76±5.01) cells (Fig. 3c,d), suggesting that *CDC20*^+^ cells are closer to completion of cell division. Not only was there a significant difference in cleavage furrow formation but the number of cells in the G2/M-*CDC20*^+^ population were higher. Based on cell counting of live cells at 1h post-sort, G2/M-*CDC20*^+^ cells were about 1.6x more numerous vs G1 and G2/M cells and about 1.3x more numerous than the G2/M-*CDC20*^−^ cell population which also had observed cleavage furrow formation (Fig. 3e).

Flow cytometry and live cell imaging confirmed the significant differences evident at 1h in overnight tracking of cellular proliferation. We used an intracellular dye (CTY) to quantify cell division by FC. With each cell division, the dye concentration dilutes due to sharing the intracellular components with daughter cells and the subsequent synthesis of new cytoplasmic components. To control for loss of signal due to intrinsic effects (described above), we normalized the fluorescence intensity observed in the G2/M populations (both *CDC20*^−^ and *CDC20*^+^) to that observed in the G1 phase. Decrease of fluorescence signal was observed in nearly the entirety of the G2/M-*CDC20*^+^ population indicating that these cells had undergone at least one cell division (Fig 4a). In contrast, progression through mitosis in the G2/M-*CDC20*^−^ population took longer, and not all these cells completed cell division during our window of observation. G2/M-*CDC20*^+^ cells had an increased percentage of dividing cells at 1, 2, 4, 6, and 18h post sort when compared to G2/M-*CDC20*^−^ with the greatest increase in percent divided at 2h (24.9±4.20) vs. (13.3±5.37) (p < 0.05, t-test) (Fig. 4b). As expected, the percentage of dividing cells declined over time as the pool of parent cells made the switch from M to G1.

**Figure 4.**
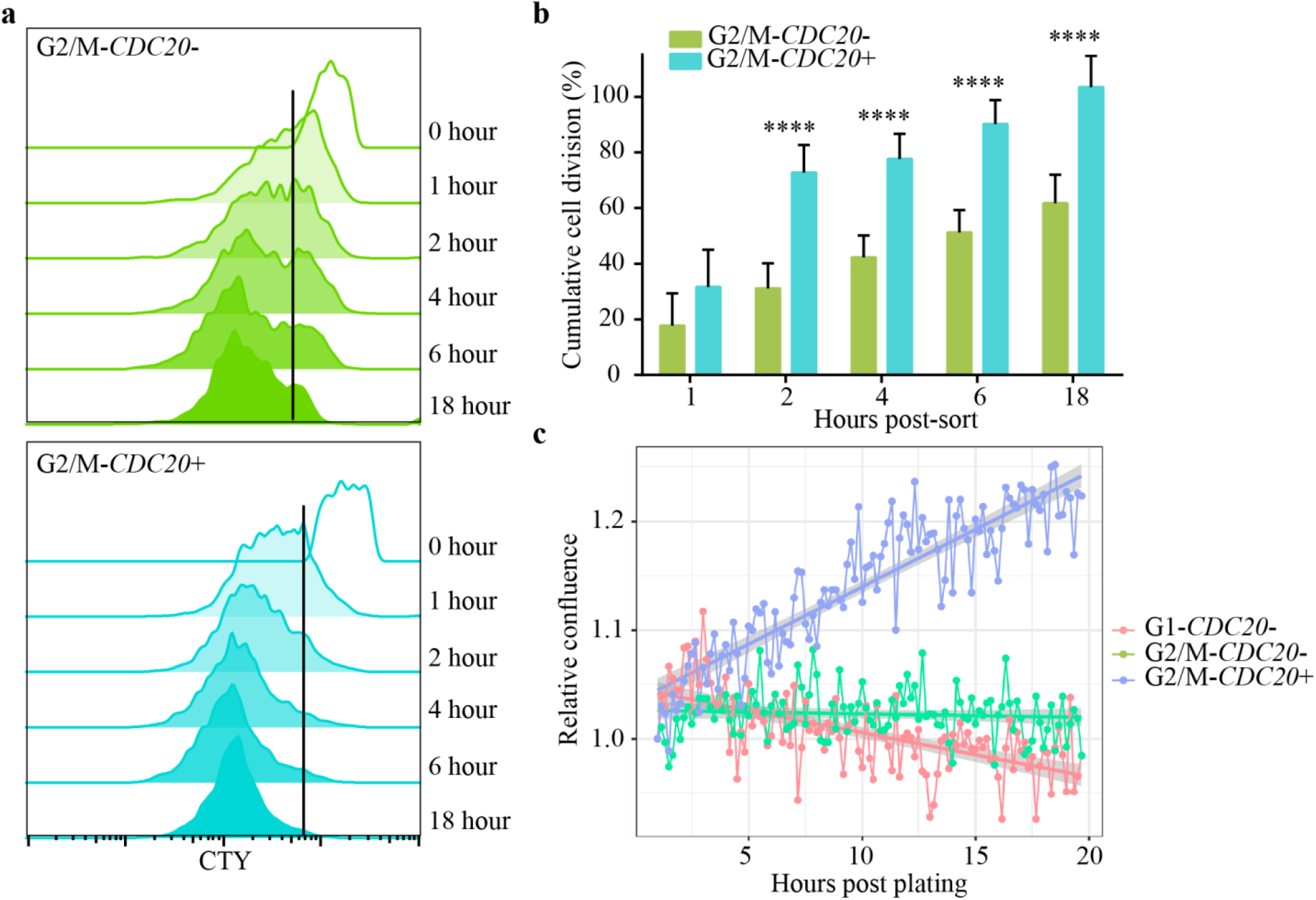
Proliferative advantage of G2/M-*CDC20*^+^ sorted events. **a)**Representative flow cytometric analysis of the proliferative ability of CellTrace Yellow (CTY) labelled D8 hiPSC-CM sorted populations, *G2/M-CDC20*^−^ (green) and G2/M-*CDC20*^+^ (blue) measured by CTY dilution after 1, 2, 4, 6, and 18h post sort. Black lines indicate fluorescence intensity at 0h, cells that divide will have fluorescence lower than the starting fluorescence. (n = 3). Due to the different starting fluorescence of parent populations at 0h, the vertical lines indicating the cells labelled with CellTrace were different. Proliferation was monitored by a decline in fluorescence relative to the starting fluorescence at 0h. **b)**Total percentage of cells as determined by FC analyses of CTY dilution from G2/M-*CDC20*^−^ (green) and G2/M-*CDC20*^+^ (blue) sorted populations. Percentages of proliferative cells were normalized to G1 percentages (set to 0). (t-test: Cumulative mean±SD; ****p < 0.0001; n = 3) **c)**Quantification of confluency occupied by cells as measured by IncuCyte™ from G1-*CDC20*^−^ (orange), G2/M-*CDC20*^−^ (green) and G2/M-*CDC20*^+^ (blue) sorted events after plating for 18h. Values represent percentage of confluency from 4 images per well normalized to 1h post-plating (set to 1). Confluency lines were fit with linear regression (population color) as 95% confidence intervals (gray). A test of the significance of the difference in slope between the regression lines was highly significant (p < 0.0001).

G2/M-*CDC20*^+^ cells had increased percentage of cells with a cleavage furrow, cell numbers, and percentage of divided cells, therefore we expected confluency to increase over time. To complement the cell division measurements by FC, we quantified proliferation by cell confluence over time in cultures of each population of interest using live cell imaging. The G2/M-*CDC20*^+^ cells exhibited increasing confluence over the 18h incubation (Fig. 4c) while the confluency of the G2/M-*CDC20*^−^ cell population was essentially flat. The decrease in G1 confluency over time suggests that cell loss was occurring in these wells and there was neither sufficient time nor pools of cells in late M phase capable of masking the cell loss. With this in mind the change in confluency in G2/M-*CDC20*^+^ cultures, and to a lesser extent G2/M-*CDC20*^−^, was even more profound suggesting that there was enough cell division to overcome decreases in viability. The difference in slope between the regression lines was highly significant (p < 0.001).

### Single-cell RNA-seq data revealed the significance of enrichment for late-mitosis CMs using MBs

To complement the functional validity for the selection of cytokinetic cells as detailed above we decided to evaluate the cells by a mechanistic approach analyzing the gene expression pattern. From the cells that were sorted by the CM surface marker SIRPA.^43^ Cells of CM lineage were identified by expression of *TMEM88*^44^ and/or *TNNT2*, thereby including CMs from both progenitor and more mature stages. The most immediate feature revealed by tSNE was a gene expression continuum for each sorted event, which correlated with cell cycle progressions (**Online** Fig. 3). Unsupervised clustering of these cells revealed significant overlap of G2/M-*CDC20*^+^ and G2/M-*CDC20*^−^ cells when clustered by genome wide expression patterns. This finding highlighted the importance of further analysis to clarify any differences between these cells in terms of cell cycle progression, as well as to identify additional markers distinguishing truly mitotic CMs.

Having sorted G2/M-*CDC20*^−^ and G2/M-*CDC20*^+^ subpopulations based on MB signal, we next examined whether the **CDC20*^+^* population was enriched for cells with gene expression patterns consistent with late mitosis. We performed scRNA-seq of cells from three populations (G1-*CDC20*^−^, G2/M-*CDC20*^−^, G2/M-*CDC20*^+^). We quantified the enrichment of cell cycle stage specific gene sets^45,46^ using a published algorithm tailored to scRNA-seq.^35^ Both the G2/M populations (*CDC20*^+^ and *CDC20*^−^) showed significant enrichment of the G2/M and M gene signature relative to the G1 population (Fig. 5a). Intriguingly, the *CDC20*^+^ population showed a significant shift in enrichment of the M/G1 gene signature, suggesting these cells were transitioning out of mitosis.

**Figure 5.**
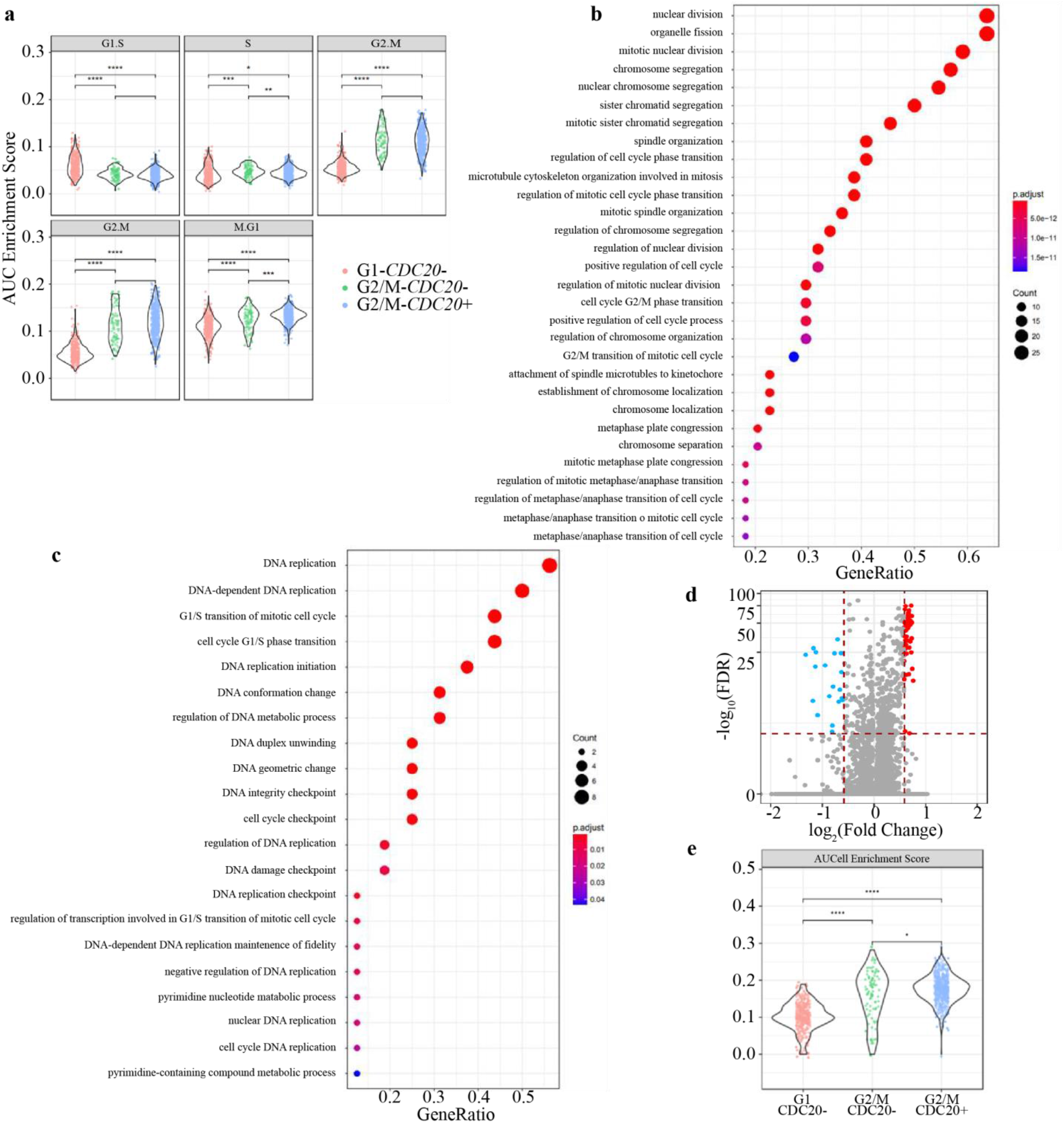
scRNA-seq reveals the *CDC20*^+^ subset of D8 hiPSC-CMs has a gene signature of mitosis progression and cytokinesis. **a)**Violin plots of cells filtered for TNNT2 > 0 or TMEM88 > 0 showing AUC enrichment scores for cell cycle stage specific gene sets from sorted *G1-CDC20*^−^ (orange), G2/M-*CDC20*^−^ (green) and G2/M-*CDC20*^+^ (blue) populations; each dot represents an individual cell. (t-test: Mean AUC score *p < 0.05 **p < 0.01 ***p < 0.001 **** p < 0.0001) **b)** and **c)**Gene Ontology enrichment analysis of the most differentially expressed (upregulated >1.5 and down regulated <1.5 fold with FDR < 0.001) genes for G2/M-*CDC20*^+^ The dot plots indicate most significant GO terms. The dotplot shows the number of genes associated with the GO terms (size) and the p-adjusted values for these terms (color). The x-axis represents different gene clusters for a single enrichment result; gene count: number of identified genes in each category; gene ratio: number of genes related to GO term / total number of signature GO genes. **d)**A volcano plot of differentially expressed genes between cell populations based on single cell expression levels; red dots indicate significantly down regulated genes (fold change > 1.5 down and FDR < 0.001) and blue dots indicate similarly upregulated genes. **e)**Violin plots showing AUCell analysis of established gene sets of components associated with cytokinesis from sorted G1-*CDC20*^−^ (orange), G2/M-*CDC20*^−^ (green) and G2/M-*CDC20*^+^ (blue) populations; each dot represents an individual cell. (t-test: Mean AUC score *p < 0.05 ****p < 0.0001).

To glean additional functional insight from the gene expression patterns in these cells, we identified those genes that were most consistently and strongly upregulated in the G1/M-*CDC20*^+^ cells relative to the other populations (**Online Table 3; Online** Fig. 4). We then performed Gene Ontology^47,48^ enrichment analysis^37^ of the most differentially expressed genes to investigate the indicated cellular role (Fig. 5d). We queried the entirety of the Gene Ontology Biological Process ontology and without exception the most enriched terms were related to nuclear division, spindle organization, and chromosome segregation (Fig. 5b). The majority of these upregulated terms specifically related to separation of the sister chromatids, whereas the downregulated differentially expressed genes were significantly enriched in DNA replication (Fig. 5c). Previous genetic and molecular studies^49–52^ have identified conserved components of the molecular machinery required to achieve cytokinesis. Our AUCell analysis of our scRNA-seq data sets revealed that this core cytokinesis gene set is enriched within G2/M-*CDC20*^+^ cells relative to the G2/M-*CDC20*^−^ population (Fig. 5e).

For the dual *CDC20/SPG20* MB enrichment strategy, AUCell analysis was conducted using the same gene signatures reflecting five phases of the cell cycle as previously characterized.^45,46^ Although there was no difference in mean AUC Score for the M.G1 signature between *CDC20*^+^*SPG20*^−^ and *CDC20*^+^*SPG20*^+^ cell populations, the *SPG20*^+^ population had a smaller standard deviation (p < 0.0001 F-test) which suggested that the *CDC20*^+^*SPG20*^+^ was a more homogenous population (Fig. 6f). Based on the genes associated with the completion of cytokinesis,^49^ AUCell analysis showed that the *CDC20*^+^*SPG20*^+^ events have a more active signature compared to other sorted events (p = 0.004, Fig. 6g).

**Figure 6.**
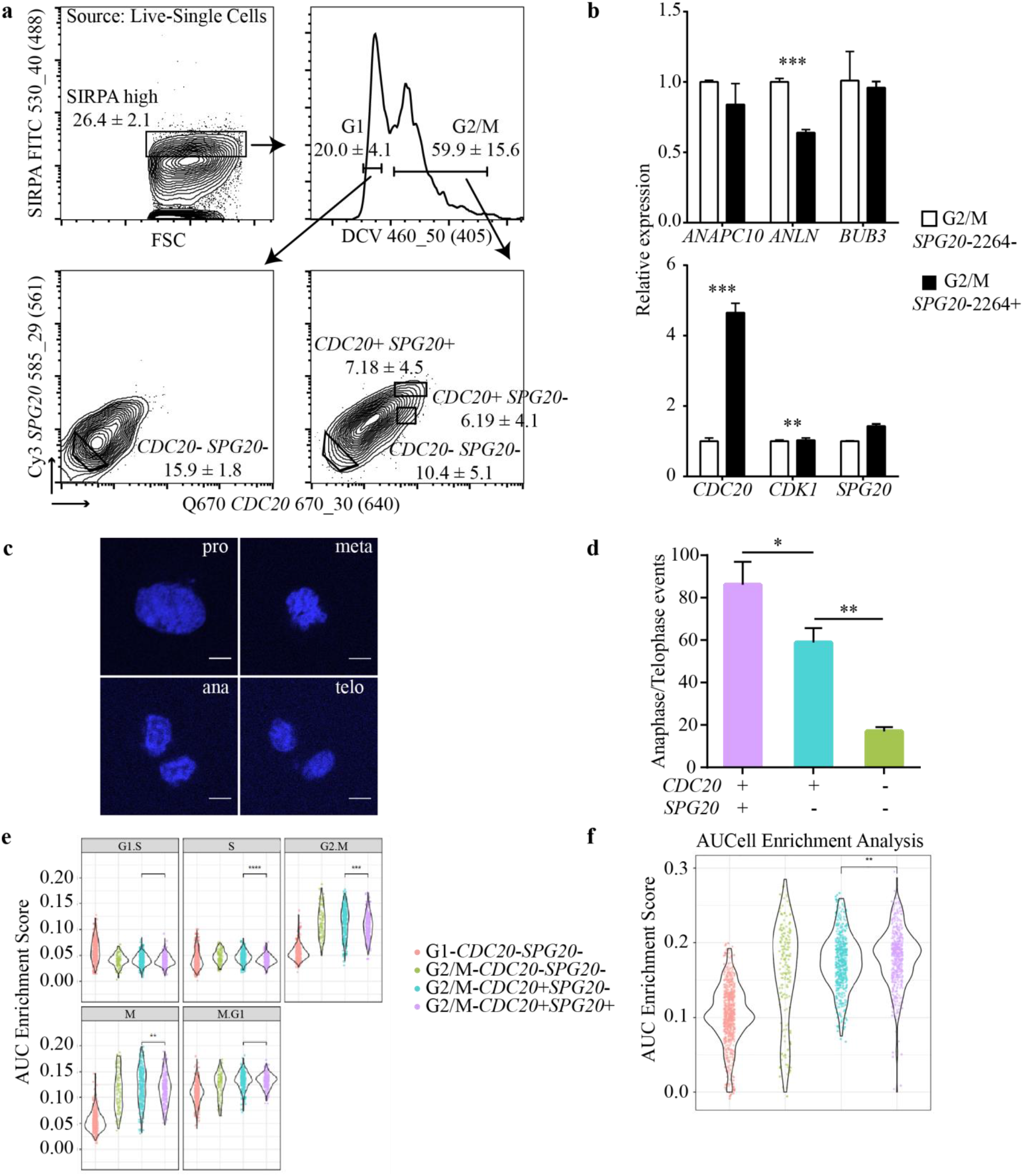
Identification and characterization of *CDC20*^+^*SPG20*^+^ D8 hiPSC-CMs. **a)**Representative FC analyses of the cell sorting strategy used for the isolation of G1-*CDC20*^−^*SPG20*^−^, G2/M-*CDC20*^−^*SPG20*^−^, G2/M-*CDC20*^+^*SPG20*^−^ and G2/M-*CDC20*^+^*SPG20*^+^ events following delivery of *CDC20* and *SPG20* MBs. Gating strategy; Debris, doublets, and non-viable events were excluded, SIRPA^high^, DCV DNA content for G1 and G2/M, *CDC20*^−^ *SPG20*^−^, *CDC20*^+^*SPG20*^−^ and *CDC20*^+^*SPG20*^+^ events. Values represent mean of parental gate percentages ± SD (n = 3). **b)**RT-qPCR analysis of the expression levels of mitosis, cell cycle, and late-M phase genes from 50 G2/M-*SPG20*^−^ (open bar) and G2/M-*SPG20*^+^ (black bar) sorted events (t-test: Mean ± SD; *p < 0.05 **p < 0.01 ***p < 0.001). **c)**Representative images of single cells counterstained with DAPI to determine the nuclear dynamics and cell cycle phase of each single cell. pro=prophase, meta=metaphase, ana=anaphase, telo=telophase; Scale bars, 20 μm. **d)**Quantification of percentages of events with late M phase nuclei morphology (anaphase and telophase) from cell sorted populations *G2/M-CDC20*^−^*SPG20*^−^ (green), *G2/M-CDC20*^+^*SPG20*^−^ (blue) and *G2/M-CDC20*^+^*SPG20*^+^ (purple). (t-test: Mean±SD; *p < 0.05 **p < 0.01). **e)**Violin plots of scRNA-seq data of sorted populations, G1-*CDC20*^−^*SPG20*^−^ (orange), *G2/M-CDC20*^−^*SPG20*^−^ (green), *G2/M-CDC20*^+^*SPG20*^−^ (blue) and *G2/M-CDC20*^+^*SPG20*^+^ (purple) comparing the AUC scores from single cells. (t-test: Mean AUC score **p < 0.01 ***p < 0.001 ****p < 0.0001). As measured by F-test the SD of the M.G1 gene signature of the *G2/M-CDC20*^+^*SPG20*^+^ cell population was significantly smaller compared to G2/M-*CDC20*^+^*SPG20*^−^ (0.0158 vs. 0.0123 p < 0.0001). **f)**AUC values for the four sorted events. The gene signature is based on components necessary for completion of cytokinesis, such as microtubule associated protein, centralspindlin complex, Aurora kinase complex, and the RhoA pathway ^59^.

### Temporal distribution of DNA dye for identification of cell-cycle stage after dual MB-based FC sorting

We investigated whether our *CDC20* enrichment strategy enhanced the percentage of cytokinetic events when combined with MBs targeting *SPG20* mRNAs. Since our cell model of D8 progenitor/committed hiPSC-CMs was unlikely to have multinucleated cells,^53^ quantification of anaphase/telophase events would provide evidence of cytokinesis. G2/M-*CDC20*^−^ *SPG20*^−^, G2/M-*CDC20*^+^*SPG20*^−^, and G2/M-*CDC20*^+^*SPG20*^+^ cell populations were individually sorted into 96-well plates and observed by confocal microscopy (Online Fig. 5). All four phases of mitosis, pro-, meta-, ana-, and telophase were observed (Fig. 6c). Microscopic observation at high magnification (100X) clearly indicated that cells undergoing karyokinesis were enriched in both G2/M-*CDC20*^+^ populations with the difference more pronounced in *G2/M-CDC20*^+^*SPG20*^+^ cells (Fig. 6a,c,d; Online Fig. 5). The number of late mitotic events increased by threefold in cells sorted based on *CDC20* MB (58.9±6.8) and four fold among cells sorted based on *CDC20/SPG20* MBs (86.1±10.8) compared to G2/M positive MB negative cells (17.0±2.0) (Fig. 6d). This demonstrates that that our flow cytometric strategy of additionally targeting *SPG20* improved our ability to isolate hiPSC-CMs undergoing cytokinesis.

### Distinguishing CM division from polyploidy and binucleation

Polyploidy and multinucleation are characteristic features of mammalian CMs. Multinucleation occurs in the last step of cell division as cytokinesis ceases. One proposed reason for cytokinesis failure during CM binucleation is a defective contractile ring.^54–57^ The contractile actomyosin ring is formed during CM binucleation, which constricts the central spindle resulting in midbody formation.^57–59^ While SPG20 plays a role in midbody abscission^28^, we tested whether our dual-MB strategy was capable of distinguishing cell divisions from binucleated events. P3 rat-CMs were treated with serum and an increase in the percentage of binucleated CMs was observed (Fig. 7a-b). On D3, ~67±6% of CMs were binucleated (Fig. 7b). Following serum stimulation, we found that both late-M phase genes *Spg20* and *Spast* were upregulated. Interestingly, *Cdc20* was down-regulated (Fig. 7c). Due to the crucial role of CDC20 at the mitotic checkpoint, the downregulation of *Cdc20* was expected among the binucleated events. It seems reasonable to conclude that while SPG20 expression may be involved in binucleation (Fig. 7d), CDC20 serves as a checkpoint for cytokinesis and can distinguish cytokinetic from acytokinetic (polyploi d/multinucleated) events.

**Figure 7.**
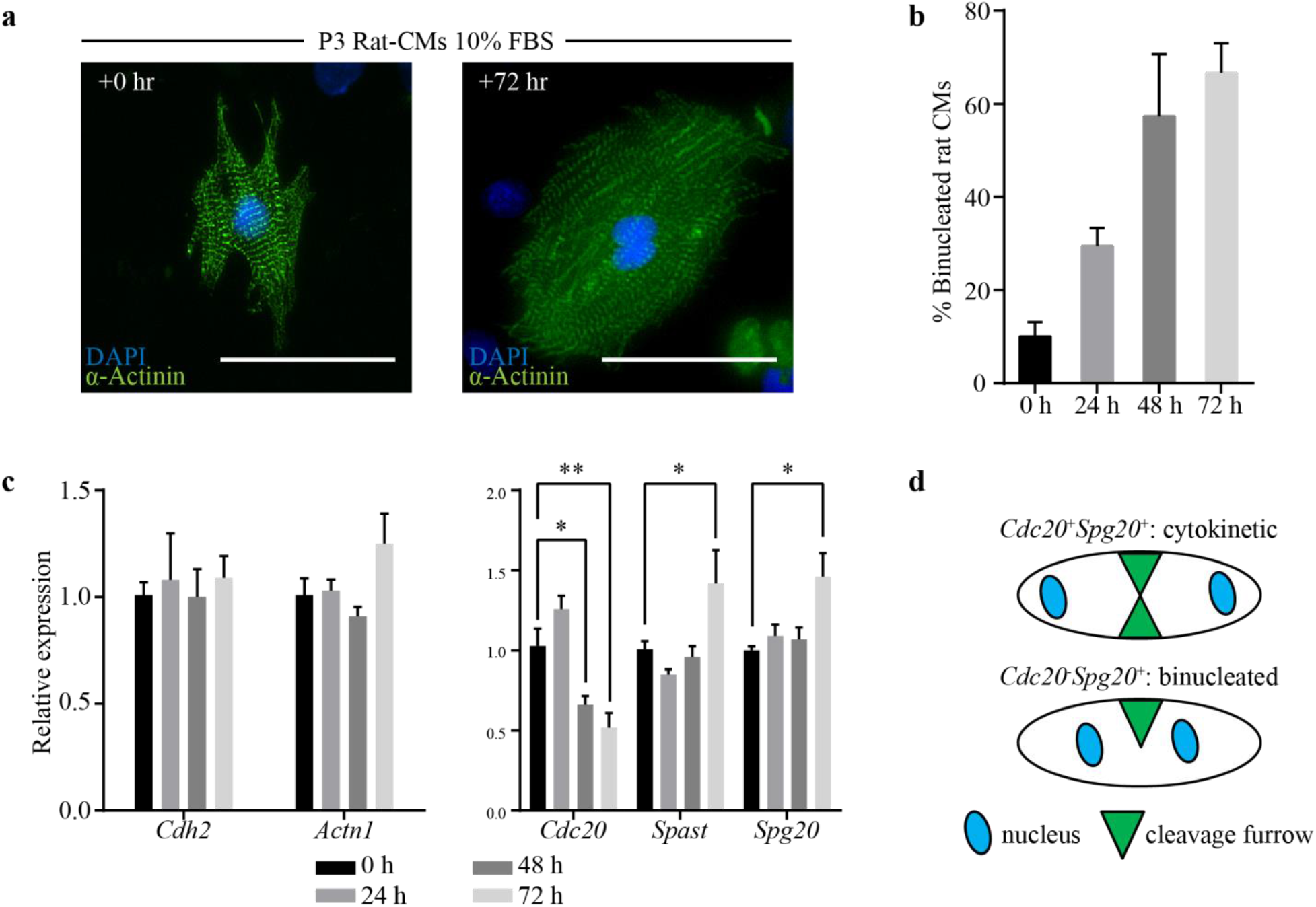
Gene expressions in binucleated rat-CMs showed decrease in *Cdc20* and increase in *Spg20*. **a)**Images showing immunostaining of structural protein α-Actinin and DAPI counterstained nuclei in P3 Rat-CMs treated with 10% FBS as binucleation progresses. Scale bar, 100 μm. **b)**Quantification of binucleated Rat-CMs treated with 10% FBS (Mean±SEM). **c)**RT-qPCR analysis of the expression levels of CM and late-M phase genes from serum treated Rat-CMs at 0 (black), 24 (gray), 48 (dark gray), and 72 (light gray) hours (t-test: Mean ± SEM; *p < 0.05 **p < 0.01). The Ct values were normalized according to the ΔΔCt method using *Ubc, Ppia*, and *Ywhaz* as the endogenous control genes and tested for significance by t-test. d)Proposed hypothesis of applying MB-based detection to distinguish cytokinetic (*Cdc20*^+^*Spg20*^+^) from binucleated (*Cdc20*^−^*Spg20*^+^) events. Schematic modified from Kadow et al, 2018.

To test the hypothesis that the expression levels of *SPG20* and *CDC20* could identify polyploid and binucleated events in the human system, we returned to hiPSC-CMs at late-stage differentiation. D100 hiPSC-CMs developed polyploidy and binucleation, resembling *in vivo* human CMs (Fig. 8a,b). D100 post *in vitro* differentiation, ~24±4% of CMs were binucleated and ~16±1% of CMs were calculated as poly(tetra)-ploid (Fig. 8b). We sorted G2/M cells in D100 cultures and checked the gene expressions of *CDC20, SPG20*, and *SPAST*. Compared to G2/M phase cells at D8 (Fig. 3a), the degree of *CDC20*-MB signal shift was completely absent (Fig. 8c). RT-qPCR results suggested that *CDC20* downregulation at D100 may be associated with the lack of cytokinesis (Fig. 8d). The lack of *SPG20* or *SPAST* differential expression at the late stage may be explained by the heterogeneous populations mixed with mononucleated (polyploid) and binucleated events. Additional analyses are required to understand the role of SPG20 during polyploidization and binucleation. Initial sorting using our dual-MB strategy on late-stage hiPSC-CMs offers a promising option for the enrichment of binucleated (*CDC20*^−^*SPG20*^+^) and polyploid (*CDC20*^−^*SPG20*^−^) events (Fig. 8d).

**Figure 8.**
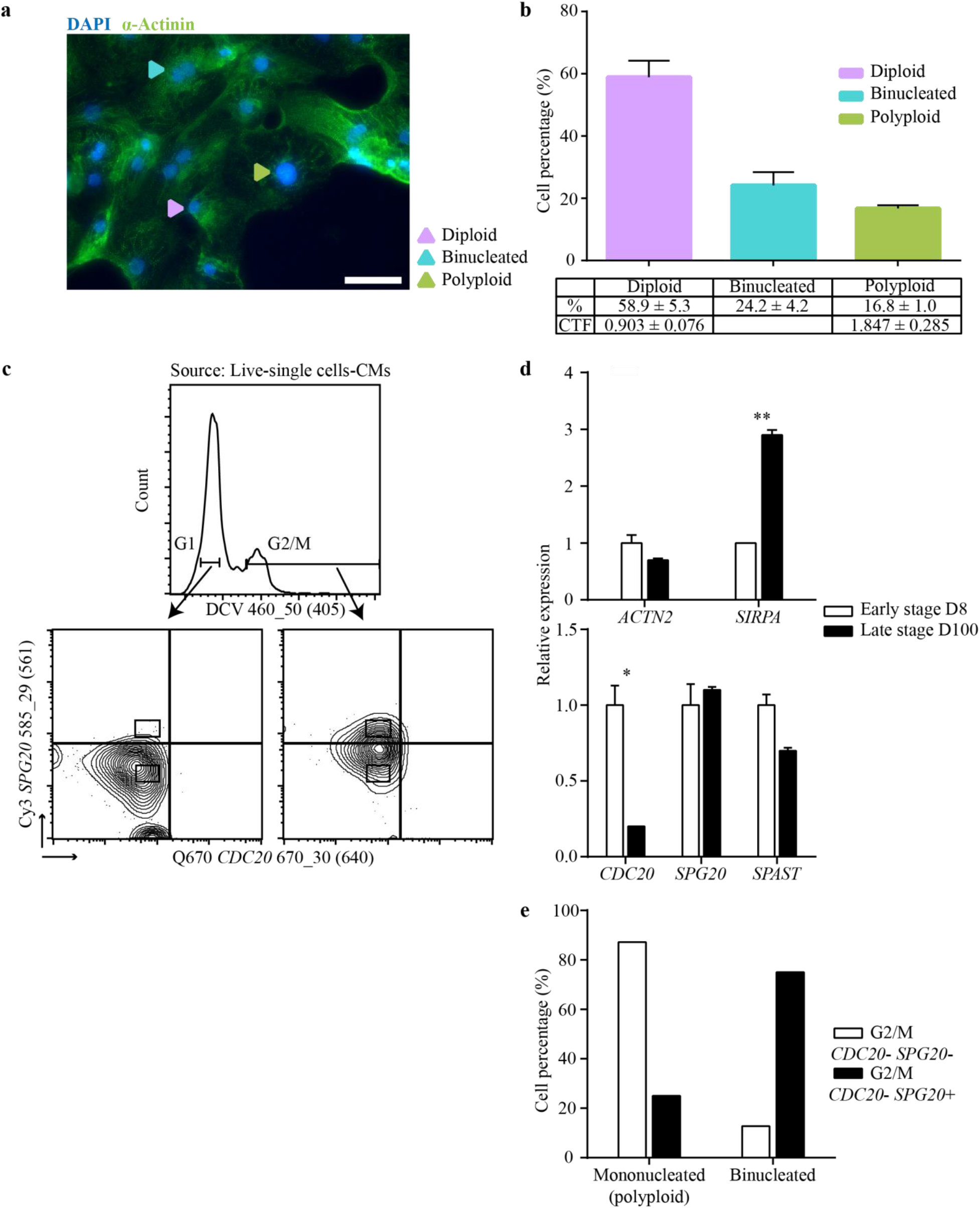
Nuclei dynamics and mRNA levels of *CDC20* and *SPG20* in late-stage hiPSC-CMs. **a)**Image showing immunostaining of structural protein α-Actinin and DAPI counterstained nuclei in hiPSC-CMs at D100. Purple arrows indicate mononucleated diploid hiPSC-CMs. Blue arrows indicate binucleated hiPSC-CMs. Green arrows indicate mononucleated polyploid hiPSC-CMs. Scale bar, 20μm. **b)**Cell percentages of diploid (purple), binucleated (blue), and polyploid (green) D100 hiPSC-CMs quantified from IF staining using ImageJ software. CTF (corrected total fluorescence) = Integrated density of selected nuclei – (Area of selected nuclei * Mean fluorescence of background readings). **c)**Sorting strategy for isolating G2/M-*CDC20*^−^ *SPG20*^−^ and G2/M-*CDC20*^+^ *SPG20*^−^ events. Gating strategy: Debris, doublets, and non-viable events were excluded, SIRPA^high^, DCV DNA content for G2/M, and *CDC20/SPG20* MB signal shifts. **d)**RT-qPCR analysis comparing the expression levels of CM and late-M phase genes from 50 sorted G2/M events using D8 hiPSC-CMs (open bar) and D100 hiPSC-CMs (black bar) (t-test: Mean±SEM; *p < 0.05 **p < 0.01). **e)**Percentage of mononucleated (polyploid) and binucleated cells determined by post-sort fluorescence microscopy.

## Discussion

Postnatally, CMs respond to the increased hemodynamic burden with an increase in cardiac mass through hypertrophy rather than hyperplasia.^9,27,60,61^ While hypertrophy is advantageous in terms of maintaining contractility and conserving metabolic resources,^62,63^ proliferative potential is compromised and prevents full recovery from injury. Following injury, interstitial fibrosis and hypertrophy occur together with permanent loss of CMs, leading to pathological remodeling, dysfunction and heart failure.^64,65^ In this study we aim to enhance our ability to study the regulation of CM proliferation towards the future goal of increasing myocardial regeneration following injury.

Since the initial reports of cardiomyocyte renewal in the adult human heart,^7,8^ the source of these new cells has remained debated. Ample evidence has been accumulated that new CMs arise from pre-existing CMs rather than adult cardiac stem cells.^13,66^ CM shape and cytoplasmic contents complicate matters—mature and functionally active CMs are elongated and are filled with mitochondria and filaments of the contractile apparatus. This has led investigators to conclude that CMs are most-likely terminally differentiated and incapable of cellular division. Nevertheless, mitosis of CMs has been observed and it has been established that differentiated/contracting CMs complete cytokinesis. This completion of cytokinesis is precluded by the disassembly of the contractile apparatus.^67,68^ With CMs requiring disassembly of their contractile apparatus to undergo mitosis the process of cell division has to be strictly regulated. The molecular events that mediate CM mitotic exit are poorly understood and in order to elucidate the underlying mechanisms resulting in activation of the endogenous cardiac regeneration process, a method for identifying CMs with true proliferative potential is needed. Careful studies of the molecular biology of these cells will be necessary to develop new pharmacological strategies to promote regeneration in the injured myocardium.

Here, we report a high throughput method for separating cytokinetic CMs from polyploidization or binucleation events. Through a combination of FC, gene expression, time-lapse microscopy, and nuclear morphology, we demonstrated that a MB based approach targeting *CDC20* and/or *SPG20* identified a proliferative subset of CMs derived from hiPSC cells. To verify cell activities and stages we used gene sets previously defined to characterize sorted events.^34,45,49,69^ Gene set analyses confirmed MB positive cells to be in late-stages of cytokinesis, including an expression pattern exit to G1 from M.

CMs derived from hiPSCs contract in culture, proliferate early in differentiation and transition into CMs with binucleated and polyploid nuclei later in differentiation. Therefore, hiPSC-CMs are a representative model system with similar cell cycle progression resembling *in vivo* CMs. hiPSC-CMs early in differentiation resemble fetal or immature CMs^70^ and long-term culturing yields hiPSC-CMs that closely resemble adult CMs^53^. Molecular profiling comparing cytokinetic and acytokinetic CMs at sequential stages of hiPSC-CMs during cardiac differentiation will provide a more comprehensive understanding of the mechanisms of cytokinetic CMs^53^. Here, we show that hiPSC-CMs transition from cytokinetic to acytokinetic as differentiation progressed from D8 to D100 (Fig. 8).

Compared to transgenic systems and fate-mapping experiments to monitor and quantify cytokinesis, ^71–74^ our approach provides a versatile means for live intracellular labeling and sorting of cytokinetic CMs without genetic perturbations. The use of our methodology opens the way to investigate downstream use and study in *in vivo* applications. In contrast to indirect assays such as EdU, Ki-67, pHH3, or Aurora B-kinase which are limited to identifying DNA-synthesis or mitosis without cytokinesis,^75,76^ the approach described here identifies CMs that are destined to complete cytokinesis and divide.

Our data suggests that binucleation with the lack of sustained *CDC20* expression appears to halt CMs at the binucleated stage. Further delineation of the continuous cell cycle trajectory and the uncoupling of cytokinesis from karyokinesis require a systematic molecular classification at the transcriptome level. The scRNA-seq approach has revolutionized the possibilities for defining distinct cellular states and diversity,^77–79^ thus further exploration of temporal changes in gene expression during mitosis could elucidate the transition from proliferation to a terminally differentiated binucleated phenotype.

In conclusion, we report a MB-based method to identify cytokinetic CMs derived from hiPSC-CMs via fluorescence-activated cell sorting based on *CDC20* or *SPG20* mRNA expressions. This study demonstrates a novel alternative to existing techniques dependent upon DNA content. Our methodology improves current strategies for identification of truly mitotic CMs by circumventing the unique challenges of cells that become polyploid or binucleated due to cytokinesis failure. Our technique will allow for further characterization of proliferative CMs and allow elucidation of the underlying mechanisms responsible for activating endogenous cardiomyocyte renewal, thereby opening the way to development of new therapeutic strategies for patients with myocardial injury.

## Acknowledgements

We thank members from the Grohar laboratory for use of the IncuCyte system and image analysis. We also thank the Van Andel Research Institute Genomics and Optical Imaging Core facilities.

