## Supplementary material for "Isolation of cardiomyocytes undergoing mitosis with complete cytokinesis": Online supplement

### Detailed Methods

#### Cell culture and CM differentiation

Episomal human iPSCs [1, 2] (Thermo Fisher Scientific) were maintained as a monolayer on Matrigel (Corning Life Sciences) coated plates and supplemented with Essential 8 Medium (Thermo Fisher Scientific). Identity and quality of the line was maintained/ensured by passage at three-day intervals and cryopreservation. Immunocytochemical characterization was carried out using antibodies against TRA-1-60, SOX2, OCT 3/4, and SSEA-4 to confirm pluripotency *in vitro*. Differentiations were conducted using hiPSCs within 15 passages from the last time the cells were thawed. For small molecule-based differentiations, hiPSCs were supplemented with RPMI 1640 (Gibco) medium containing B27 minus insulin (RPMI/B27 -insulin) with 6 µM of the GSK3 inhibitor CHIR990216 at D0-1 to induce mesoderm, followed with cardiac differentiation at D4-5 by changing media to RPMI 1640/B27 (-insulin) with 5 µM Wnt inhibitor IWR1 [2]. Typically, spontaneous contraction in cultures was observed at D7-8 of differentiation. Cells were harvested at D8 (progenitor/committed CMs) for flow cytometry (**Fig. 1a**).

#### Purification of human CMs through glucose starvation and long-term maintenance of hiPSC-CMs

From day 10-13 post-differentiation, cells were treated with RPMI 1640 Medium (-L-glutamine and -glucose) (Biological Industries USA, Inc.) containing B27 with insulin (RPMI/B27 +insulin). At day 13, cells were returned to RPMI/B27 +insulin and stabilized for 48 h. Cells were then dissociated into single cells with TrypLE Select 10X (Gibco) for 5 min at 37°C, neutralized with RPMI/B27 +insulin, and centrifuged for 5 min at 100 xg. Cells were then re-suspended in RPMI/B27 +insulin with 10% FBS and replated onto a new Matrigel-coated well at a seeding density of 2 million cells per well of a 6-well dish to ensure high cell survival. After 24 h, 10% FBS was removed by changing the medium back to RPMI/B27 +insulin. For long-term hiPSC-CMs maintenance, media was replaced with fresh, pre-warmed RPMI/B27 +insulin every three days.

#### Design and delivery of MBs into hiPSC-CMs

mRNA sequences of target genes were retrieved from NCBI (FASTA preferred, RefSeq prefixed with NM__) and the secondary structure of the mRNA was predicted using the online mFOLD server [3, 4] (http://unafold.rna.albany.edu/?q=mfold). Different stem designs (length and sequence) were compared based on the predicted hairpin structures using the Quickfold server [5, 6] (http://unafold.rna.albany.edu/?q=DINAMelt/Quickfold) or IDT OligoAnalyzer 3.1 (https://www.idtdna.com/calc/analyzer). MBs were designed to have high target affinity with predicted melting temperatures that prevent interference during downstream product amplification. The MB contained a fluorophore (6-FAM, Cy3, TAMRA, or Quasar 670) and a quencher molecule (Black Hole Quencher 2) at the 5' and 3' ends of the stem-loop hairpin structure. To achieve high specificity, the target sequences of MBs were analyzed using BLAST; only MBs returning E-values less than 1 for the target gene and greater than 1 for other genes were selected for synthesis. A negative control random probe (negative control MB) was designed (**Supplementary Table 1**) with a 17-base probe sequence with no complimentary match in the mammalian genome, preventing hybridization and fluorescence from subsequent unfolding. This negative control MB was designed with both a quencher and a fluorophore in order to allow monitoring of background fluorescence, enzymatic degradation, and serve as a flow cytometry gating control (**Fig. 1b,d**). In contrast, a positive control MB with a single fluorophore (no quencher) attached to the 5' end of the stem was designed (**Supplementary Table 1**) to serve as a compensation control and indicator of delivery efficiency. In order to facilitate cellular delivery, all MBs were modified with the cell-penetrating peptide MPG (GALFLGFLGAAGSTMGAWSQPKSKRKV, with N-terminal acetylation and C-terminal cysteamide modification) to form a stable non-covalent complex and promote the uniform and rapid delivery of MBs into cells without interfering with specific targets or hybridization-induced fluorescence [7, 8]. All HPLC (high performance liquid chromatography) purified MBs were ordered from GenScript and LGC Biosearch Technologies and MPG was ordered from GenScript. The MB-MPG complexes were formed in 500 µl of RPMI/B27 +insulin and incubated with gentle agitation (orbital shaker 50 rpm) for a minimum of 1 h (up to 1.5 h) in a standard 37°C 5% $CO_2$ incubator. MBs were incubated with MPG (MPG charge +4) at a molar charge ratio of 1:25. For every $1x10^6$ cells in suspension, 1 µl x 10 µM of MB concentration was added. After 1 h incubation at 37°C on an orbital shaker, $1x10^6$ cells were added to the MB/MPG complex medium. Additional growth medium was added to bring the final volume to 1 ml per well when working with 24-well plates (Nest). (Protocol optimization based on delivery efficiency and cell viability should be established prior to working with other cell culture plates with different combinations of format and surface). Following MB/MPG complex formation additional labeling reagents were added according to the assay being performed and returned to the 37°C incubator for 1 h with gentle agitation (orbital shaker 50 rpm). Following the 1 h incubation, cells were washed with sorting buffer (1% FBS/0.5 mM EDTA in 1x HBSS without $Ca2^+/Mg2^+$) for 5 mins at 100 x g at room temp. Cell pellets were resuspended in sorting buffer and transferred through a 35 µm filter into a new 5 ml polypropylene round-bottom tube (Falcon), and analyzed/sorted on a BD Influx (BD Biosciences) flow cytometer using Sortware v1.2.

**Separation of live cells in different cell cycle phases**

To enrich for cells in G1 and G2/M cell cycle states, hiPSC-CMs were labelled with titrated amounts of Fixable Viability Dye eFluor 780 (fvd780) (Thermo Fisher Scientific), FITC anti-SIRPα/$\beta$ (SIRPA) [9] (Clone SE5A5 BioLegend), and Vybrant DyeCycle Violet Stain (DCV) (Thermo Fisher Scientific). Cultured differentiating hiPSC-CMs at D8 were dissociated with stem cell gentle dissociation reagent (Stemcell Technologies) and resuspended at a concentration of $1x10^6$/ml in RPMI/B27 +insulin. Staining for flow cytometry was completed at 37°C. For all experiments DNA was labelled with 5 µM DCV for 1 h, cells were labelled with 0.44 µg/ml SIRPA for 30 mins, and non-viable cells were labelled with 1 µl/ml of fvd780 for 30 mins. Labeled cells were washed twice in sorting buffer for 5 mins at 100 x g at RT and run on a BD Influx flow cytometer. Combined with the fact that this study was focused on live cell sorting and unavailability of algorithms for the deconvolution of DNA-content-frequency employed in real-time, our DNA content gating strategy was based on previous published use of the DyeCycle reagent catalog to isolate live cells in different cell cycle stages [10]. The flow sorting of hiPSC-CMs in G1 andG2/M phase of the cell cycle was defined by the following gating hierarchy: hiPSC-CMs were separated from debris based on distribution of light scatter by SSC/FSC; cell doublets were excluded by signal pulse characteristics of FSC-W/FSC-H and SSC-H/SSC-A. Viable cells with intact cell membranes were gated as fvd780 negative. Differentiated CMs were gated as $SIRPA^{high}$. The DNA profile of live hiPSC-CMs was characterized by the fluorescence intensity of DCV, gates in the DNA histogram were drawn on G1 and G2/M peaks. Fifty sorted events for both G1 and G2/M populations were sorted with a BD Influx equipped with 488, 561, 405, 355, and 640nm laser lines. Sorting was performed with a 100 µM nozzle, 20 psi, a drop frequency of 39 KHz and an offset of 1.0. Acquisition of flow cytometry data was performed on a BD Influx using Sortware v1.2 and flow plots were generated with FlowJo v10.0.8.

**Validation of MBs targeting *CDC20* or *SPG20* mRNAs**

Experimental probes were required to meet both flow and RT-qPCR checkpoints before proceeding with further experiments. For MBs complementary to human-specific *CDC20* or *SPG20* mRNAs, multiple candidate target sequences were designed for each target gene and tested for fluorescence shift prior to cell sorting for cellular amplification of mRNA targets. Probes with the highest fluorescence were considered to pass the flow QC (quality control) and subjected to the RT-qPCR checkpoint. For both *CDC20* and *SPG20* MBs, 50 events were sorted from MB positive and MB negative cell populations and the expression levels of cell cycle, cell cycle regulator, mitosis, proliferation, and late M-phase genes were compared to test our enrichment hypothesis for both probe targets. MBs with a significant upregulation, t-test of 3 independent experiments, of required genes were selected for further experiments.

**Isolation of target cells using flow based on MB signal**

To further separate cells in G1, G2 and M cell cycle phases we built upon our DNA content sorting strategy and employed a mRNA target approach on D8 hiPSC-CMs. Since exclusive markers for flow sorting of live cells in M phase are not available, we delivered MBs targeting *CDC20* or *SPG20* and sorted based on shifts in fluorescence of Q670, Cy3, or TAMRA respectively. Cultured differentiating hiPSC-CM at D8 were dissociated as described above and resuspended at $1x10^6$/ml of RPMI/B27 +insulin. MBs were delivered and cells were subsequently labelled with DCV, SIRPA, and fvd780. Following labelling, cells were washed twice with sorting buffer. The flow sorting of hiPSC-CMs in G1, G2, and M phases of the cell cycle were defined by the gating hierarchy as described above with the addition of sort gates based on molecular beacon fluorescence. In the *CDC20* single probe approach, sorting gates were placed on G1-*CDC20*$^-$, G2/M-*CDC20*$^-$ and G2/M-*CDC20*$^+$ events (**Fig. 3a**) (**Supplementary Table 2**). In the *CDC20*:*SPG20* dual MB sorting strategy, the sorted hiPSC-CMs populations were G1-*CDC20*$^-$*SPG20*$^-$, G2/M-*CDC20*$^-$*SPG20*$^-$, G2/M-*CDC20*$^+$*SPG20*$^-$, and G2/M-*CDC20*$^+$*SPG20*$^+$ (**Fig. 6a**) (**Supplementary Table 2**). To establish the best and most reproducible strategy for determining sort gates of MB fluorescence, we used a combination of gating controls for each experiment. Each MB signal was compared against a FMO control and a negative control MB. Fluorescence associated with the negative control MB was attributed to a non-specific signal and thus excluded from the positive gate. In addition to the FMO, we used a biological control of cells in G1, reasoning that mRNA copy numbers of late M specific *CDC20* (Q670) and *SPG20* (Cy3) would not be high enough in G1 cells to cause a shift in MB fluorescence, therefore positive gates were set above MB signal from G1 gated cells. In each experiment, positive gates for *CDC20* and *SPG20* were drawn above negative controls and at the highest portion within the fluorescence contour. In order to determine gates of MB negative populations, the fluorescence shift from the positive control MB was taken into consideration. Typically 90% of cells that received the positive control MB displayed a shift in fluorescence, with this in mind we placed our negative gate below the lowest distribution of this shift, reasoning that fluorescence above this level was related to mRNA detection and therefore excluded from the negative gate.

### Gene expression analysis (RT-qPCR)

To analyze gene expression, cells were dissociated to a concentration of $1x10^6$/ml and labelled as described above. In brief, cells were incubated with the appropriate MB and labelled with 5 μM DCV, 0.44 μg/ml SIRPA, and 1 μl/ml of fvd780 prior to flow cytometry sorting. Gene expression analysis was performed using the CellsDirect One-Step qRT-PCR kit (Invitrogen), which facilitates the cDNA synthesis and specific target amplification (pre-amplification) of genes of interest. Fifty events were sorted by a BD Influx flow cytometer directly into individual PCR tubes containing 2.5 μL CellsDirect 2X reaction mix (Invitrogen), 0.05 μL SUPERase RNase inhibitor (Ambion), 1.25 μL 0.2X assay mix, 0.6 μL TE buffer (Invitrogen) and 0.6 μL SuperscriptIII/Platinum Taq mix (Invitrogen). The 0.2X assay mix contained a pool of TaqMan assays (Applied Biosystems) at a 1:100 dilution of each assay in TE buffer. Cell lysis, reverse transcription and specific target pre-amplification were performed on the Applied Biosystems Veriti 96-well thermal cycler immediately after sorting as follows: 60°C for 15 mins, 95°C for 2 mins, and 20 cycles of specific target amplification (95°C for 15 seconds, 60°C for 4 mins). Complementary DNA was diluted 1:5 with TE prior to qPCR on the Bio-Rad CFX96 Touch Real-Time PCR Detection System (95°C for 30 s, 40 cycles of 95°C for 10 s, and 60°C for 30 s) using iTaq™ Universal Probes Supermix (Bio-Rad). A no RT control as well as a non-template control was also included in the reactions. The threshold cycle ($C_t$) was determined using the Bio-Rad CFX Manager software. The resulting Ct values were normalized according to the ΔΔCt method using *ACTB* and *GAPDH* as endogenous control genes and tested for significance by t-test.

### Generation of single cell GEMs and sequencing libraries

On a BD Influx, cells were sorted into collection buffer (10% FBS/0.5 mM EDTA in RPMI/B27 +insulin) at 4°C, concentrated, inspected for viability, and counted. Sorted suspensions were added to 10X Chromium RT mix aiming at encapsulating 1800 cells per experiment. Downstream cDNA synthesis (14 PCR cycles) and library preparation were carried out as instructed by the manufacturer [11] (Chromium Single Cell 3' Library & Gel Bead Kit v2). Briefly, cells from each population were loaded into a separate sample well of the microfluidics device. The device then encapsulated the cells into individual droplets containing lysis buffer, RT reagents, and barcoding bead. Reverse transcription occurred within the droplets producing barcoded cDNA from each cell, where each cDNA sequence was labeled with (1) a bead (cell) specific barcode and (2) a UMI barcode. The droplet emulsion was then broken and sequencing libraries were prepared from each sample according to the Chromium protocol. During library preparation, the cDNA from each sample was ligated to 10X Genomics sample indexes allowing multiplexing of the samples on a single flow cell. The libraries were standardized for cDNA mass, pooled, and sequenced on an Illumina NextSeq 500 sequencer using (26 bp for read 1 and 124 bp for read 2), resulting in an average depth of 200,000 reads per cell (raw) with 81% of the bases having quality scores of 30 or better. Base calling was done by Illumina NextSeq Control Software (NCS) v2.0 and output of NCS was demultiplexed and converted to FastQ format the Cell Ranger mkfastq pipeline (10X Genomics).

### Bioinformatics analysis of single cell gene expression data

Demultiplexed samples were aligned to the reference human genome (version GRCh38) and converted to mRNA molecule counts using the Cell Ranger count pipeline, which is based on the STAR aligner. Each sample was aligned and counted, and then the counts were aggregated and normalized (for sequencing depth and total counts), resulting in a single dataset of counts-per-transcript for each gene in each cell and each sample.

**Gene set enrichment**

Signatures for each phase of the cell cycle, as well as the process of cytokinesis were curated from the literature [12, 13]. Enrichment of gene signatures in scRNA-seq data requires methods that account for the high number of undetectable transcripts in each cell. Therefore, we used the AUCell algorithm [14] to perform gene set enrichment analysis on our dataset. The expression of each transcript in each cell was ranked, with random shuffling of ties including non-expressed transcript. Scoring based on ranks, eliminates *rank-invariant* biases related to expression units and normalization procedure. Scores for each cell within each population were plotted, and differences between groups with respect to the AUCell score was evaluated by t-test.

**Differential gene expression**

As with gene set enrichment analysis, identification of differentially expressed genes requires an approach which accounts for the relative sparseness of the data matrix (due to undetected transcripts). We used the pagoda2 package [15] to identify genes that were differentially expressed between cell populations based on single cell expression levels. The data set was filtered to exclude cells with less than 500 transcripts and genes with less than 10 transcripts across all cells. The data was then normalized to reduce expression level-dependent bias in variance.

**Gene ontology term enrichment**

Enrichment for gene ontology terms was calculated with the help of the DOSE package [16]. This package uses the hypergeometric distribution to estimate the probability of overlap between a gene signature of interest (in our case, gene enriched in a particular cell population as calculated above) and genes in functionally predefined signatures. As our reference signature data set we used the genes associated with each GO term as defined in the org.hs.eg.db annotation package which is in turn derived from annotations from the Gene Ontology Consortium [17] [18]. For this analysis, we used those genes that exhibited differential expression in the MB positive subpopulation relative to other populations with false discovery rate < 0.001 and a fold change of at least ± 1.5.

**Dual probe sorting nuclei sorting strategy and fluorescence microscopy**

In order to determine if our sorting strategy was able to enrich for cells in M-phase and to quantitate the cell cycle status of individual cells we sorted 96 single cells from three cell populations G2/M-$CDC20^{-}SPG20^{-}$, G2M-$CDC20^{+}SPG20^{-}$ and G2M-$CDC20^{+}SPG20^{+}$ (**Fig. 6a,c**) (**Supplementary Table 2**). Cells were suspended in sorting buffer and events were individually sorted into Matrigel-coated 96 well plates (1 cell/well) CC2 Optical CVG plate (Thermo Scientific) covered with 2.5µl of freshly prepared 1% PFA. Following fixation collected events were then covered with ProLong Diamond Antifade Mountant with DAPI (Thermo Fisher Scientific) to prevent drying and to ensure nuclei visualization. The images were acquired on a Nikon A1 Plus-RSi laser scanning confocal microscope equipped with Hamamatsu

Orca Flash 2.8 sCMOS camera. A 100X Plan Apo TIRF DIC- Oil immersion and Nikon Elements acquisition and analysis software were used to observe the mitotic structures in sorted events/wells to identify the stages of mitosis, including prophase (pro), metaphase (meta), anaphase (ana), and telophase (telo) (**Fig. 6d**). Observed mitosis stages were quantified and compared by t-test to determine significance.

**Proliferation assay – CellTrace Yellow**

In all proliferation experiments hiPSC-CMs (D8) were labelled with 6 µM CellTrace Yellow (CTY) (Thermo Fisher Scientific). CTY is a cell permeable intracellular stain characterized by high fluorescence intensity. Following dissociation cells were resuspended at $1x10^6$/ml for subsequent MB delivery and labelled with 5 µM DCV, 0.44 µg/ml SIRPA, and 1 µl/ml of fvd780 prior to flow cytometry sorting. CTY dye concentration was titrated to achieve the brightest signal and least amount of cell toxicity at 48 hours post culture. Cell sorting was performed using a BD Influx flow cytometer with the following gating hierarchy: Debris, cell doublets, and non-viable cells were excluded; differentiated CMs were gated as $SIRPA^{high}$, G1 and G2/M peaks were gated, *CDC20* positive and negative gates were drawn and finally sort gates were set on CTY fluorescence for the following cell populations G1-*CDC20*$^-$CTY$^+$, G2M-*CDC20*$^-$;CTY$^+$, and G2M-*CDC20*$^+$;CTY$^+$. Sort gates in the CTY channel were placed on the peak fluorescence of each histogram and had a CV of approximately 20. Cell sorting parameters for CTY experiments were performed with an offset of 1.0, 100µm nozzle at 20psi and 39 kHz with a sort mask of 1.0 drop pure. Sorted events (10,000) were collected in 200 µl of collection buffer and added directly to individual wells of a 96 well plate coated with Matrigel and placed in a humidified 37°C 5% $CO_2$ incubator. At specified time points, the supernatant was collected and adherent cells were treated with 100 µl of stem cell gentle dissociation reagent for 5 mins at 37°C, cells were removed with the addition of 200 µl of pre-warmed RPMI/B27 +insulin and combined with supernatant and passed through a 35 µm filter. To exclude non-viable cells from analysis, 3 nM of SYTOX Green (Thermo Fisher Scientific) was added and cell solutions were analyzed for changes in CTY fluorescence on a BD Influx. To ensure that machine related error did not contribute to a decrease in CTY signal we analyzed 8 peak beads (Spherotech) prior to each time point to confirm that laser alignment was consistent between the measured time points (**Supplementary Fig. 2a**). Decreases in CTY (dividing CMs) were determined by comparing time course histograms to the parental (0 h) histograms for sorted populations G1-*CDC20*$^-$CTY$^+$, G2M-*CDC20*$^-$;CTY$^+$, and G2M-*CDC20*$^+$;CTY$^+$. To correct for non-biologically related CTY dilution we normalized fluorescence loss by subtracting decreased fluorescence percentages of G2M-*CDC20*$^-$;CTY$^+$ and G2M-*CDC20*$^+$;CTY$^+$ populations from decreases in G1-*CDC20*$^-$CTY$^+$ percentages. Normalized percentages from 3 independent experiments were averaged and compared by t-test for significance. In order to determine if efflux of CTY dye was occurring in D8 hiPSC-CMs we treated CMs with [10ng/ml] Nocodazole for 48 hours labelled with 5 µM DCV and 6 µM CTY and sorted for live; arrested G2/M;CTY$^+$ cells (**Supplementary Fig. 2b**). Following sorting, cells were plated as described above with the exception of Nocodazole treatment throughout the remainder of the experiment. Cells were harvested and analyzed for decreases in CTY fluorescence at 1 and 2 h.

**Quantitative measurement of proliferation after cell sorting via *in vivo* PKH67 tracking**

For live imaging and proliferation quantification experiments we employed our single *CDC20* (**Fig. 3a**) and dual *CDC20/SPG20* (**Fig. 6a**) MB sorting strategy as described above. In short, cells from G1-*CDC20*$^{-}$, G2/M-*CDC20*$^{-}$, G2/M-*CDC20*$^{+}$, G1-*CDC20*$^{-}$*SPG20*$^{-}$, G2/M-*CDC20*$^{-}$*SPG20*$^{-}$, G2/M-*CDC20*$^{+}$*SPG20*$^{-}$, and G2/M-*CDC20*$^{+}$*SPG20*$^{+}$ (**Supplementary Table 2**) were sorted for live imaging. 96-well tissue culture plates (Falcon) were pre-coated with Matrigel. Sorted cells were centrifuged at 150 x g for 5 mins at RT. Cells were then labeled using PKH67 Fluorescent Cell Linker kits (Sigma-Aldrich), with minor modifications in the washing process. The cell pellets (< 25 µl of supernatant) were resuspended in 250 µl of Diluent C (cell solution). PKH67 dye (1 µl) was diluted in 250 µl of Diluent C (PKH67 solution). Then, 250 µl cell solutions and 250 µl PKH67 solution were mixed in a 2 ml centrifugation tube without vortexing. Samples were mixed gently for 2 mins and 1.5 ml of collection medium (RPMI/B27 +insulin with 10% FBS) was added to bind the excess PKH67 dye. PKH67-labeled cells were centrifuged at 150 x g for 5 mins at RT. Finally, PKH67-labeled cell pellets were resuspended in 200 µl of collection medium and transferred into the Matrigel-coated plates at a concentration of 15,000 cells/96-well. Plates were mounted on the IncuCyte Zoom imaging platform (Essen Bioscience) which was maintained at 37°C with 5% CO2. Real-time images were acquired for the same region in each well every 10 mins overnight. Phase and green fluorescence (PKH67) images were captured using a 10X objective, and a four-field scan pattern was used for each well. Cell confluence was determined and analyzed using the IncuCyte ZOOM System by applying an IncuCyte mask analyzing green-fluorescent images of each well. After carefully examining the IncuCyte Images, cell confluence was normalized to 60 mins post-plating (as approximately 60 mins was required for the sorted cells to settle onto the well). A linear model was fit to the resulting data, with dummy variables added to distinguish between the populations in each well. By including an interaction term between the population variable and the confluence score at each time point, significance of the difference in slope between the subpopulations was be calculated.

#### Assays for investigating binucleation and polyploidization using rat-CMS and late-stage hiPSC-CMs

R-CM-561 Clonetics™ Rat Cardiac Myocytes (P1-3) were purchased from Lonza. Thawing of cells and initiation of culture process were performed according to the manufacturer's directions using rat cardiac growth media without adding BrdU or FBS (R313F-500, Cell Applications, Inc.). Cells were counted and cultured for 72 h at a seeding density of 250,000 cells/12-well (Corning Life Sciences) or 25,000 cells/96-well (Nunc™ MicroWell™ 96-Well Optical-Bottom Plates with Coverglass Base, Thermo Fisher Scientific). Both 12-well and 96-well plates were pre-coated with 25 µg/ml fibronectin (150 µl for 96-wells and 500 µl for 12-wells, Sigma-Aldrich, F1141) for 1 h at 37°C. On day 3, rat-CMs were stimulated with 10% FBS using Rat Cardiomyocyte Culture Medium Kit (R313K-500, Cell Applications, Inc.) for 0, 24, 48, and 72 h. After dissociating the cells in 12-wells, a commercial kit was used (RNeasy Mini Kit, Qiagen) and the total RNA was extracted along with on-column DNase digestion according to the instructions of the manufacturer. NanoDrop™ was used to determine the concentration of RNA and 500 ng of total RNA eluate was reverse transcribed using 200 units of Superscript III RT (Invitrogen) and 500 ng of oligo(dT)$_{12-18}$ primer (Thermo Fisher Scientific). Quantitative PCR was carried out on the Bio-Rad CFX96 Touch Real-Time PCR Detection System (95°C for 30 s, 40 cycles of 95°C for 10 s, and 60°C for 30 s) using iTaq™ Universal Probes Supermix (Bio-Rad). For immunofluorescence analysis, cells in 96-wells were fixed with 4% paraformaldehyde for 10 mins at 4°C and permeabilized for 20 mins at RT with 0.1% Triton X-100 and 200 mM glycerin in PBS. Cells were then blocked for 3 h (2% BSA, 0.1% Triton X-100, and 10% donkey serum from Abcam ab7475) followed by overnight incubation with primary antibody anti-α-actinin at 4°C (1:50, Invitrogen MA1-22863) and 1 h with secondary antibodies conjugated to Alexa Fluor 488 (1:1000, Abcam, ab150117) at RT in blocking buffer. After three washes (0.1% BSA, 0.1% Triton X-100), cells were mounted in ProLong™ Diamond Antifade Mountant with DAPI (Invitrogen). High resolution images were captured on the Leica DFC3000 G system using a 40X objective (Leica N Plan L 40X/0.55) and analyzed with LAS V4.6 software.

For long-term culture of hiPSC-CMs, following glucose starvation, cells were replated onto a new Matrigel-coated well at a seeding density of 2 $x10^6$cells per 6-well dish or 50,000 cells per 96-well as described above. Cells were maintained by replacing media with fresh and pre-warmed RPMI/B27 +insulin every three days until the end of experiment on day 100. On the day of experiment, immunostainings and fluorescence microscopy were performed as described above. Primary antibody: mouse anti-α-actinin (1:50, Invitrogen MA1-22863). Secondary antibody: goat anti-mouse IgG H&L (Alexa Fluor 488) preadsorbed (1:1000, Abcam, ab150117). Images were analyzed with ImageJ software to determine ploidy level based on the integrated intensity of nuclei staining (CTF corrected total fluorescence = integrated density of selected nuclei – (area of selected nuclei * mean fluorescence of background readings)). Human iPSC-CMs in G2/M phases of the cell cycle were sorted followed by RT-qPCR to determine the mRNA level of *CDC20* and *SPG20*.

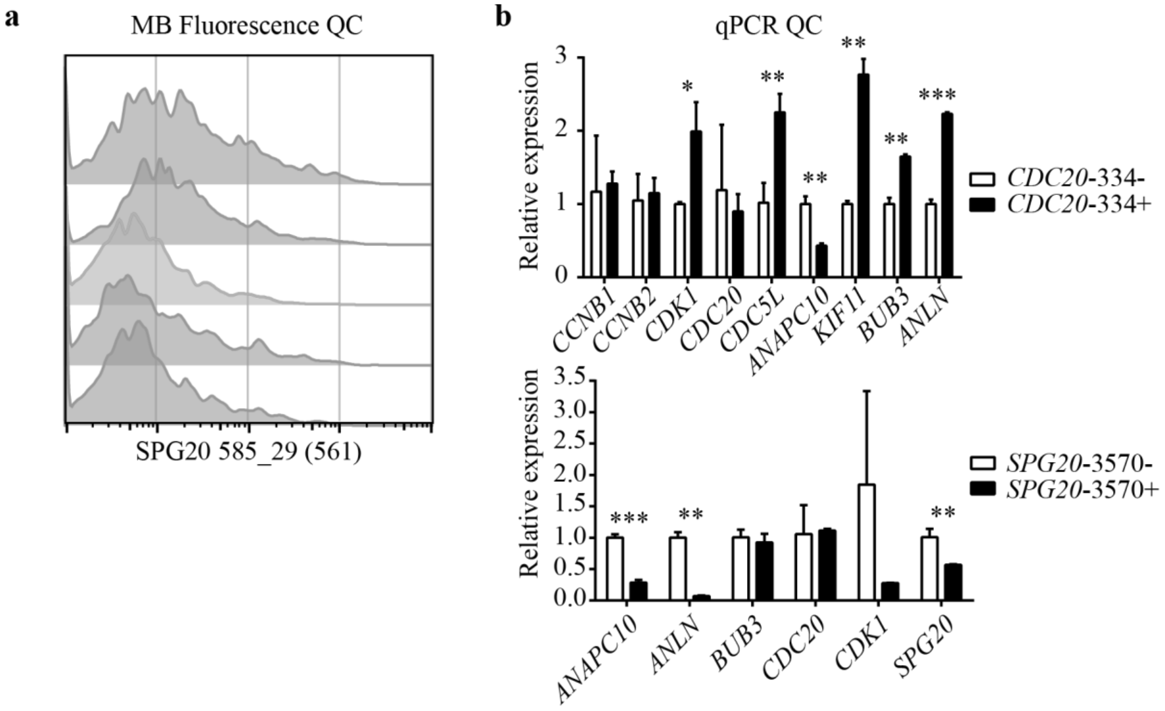


**Supplementary Figure 1. Quality control checks for molecular beacon selection. a)** Example of a failed QC step 1: Representative flow cytometry analyses of *SPG20* molecular beacon candidates. Histograms were normalized to the mode and show Cy3 fluorescence for 5 different *SPG20* probes. Based on fluorescent shift alone *SPG20*-2264 (blue) and *SPG20*-3570 (red) were selected for cell sorting and RT-qPCR verification. For flow cytometry analyses; Debris, doublets, and non-viable events were excluded, remaining events were gated on DNA (DCV) G1 and G2/M peaks and analyzed for MB signal. **b)** Example of a failed QC step two: RT-qPCR analysis of the expression levels of genes that were required to be upregulated in MB positive (black bar) vs. MB negative (open bar) sorted cells. Fifty events were sorted for each population, both probes *CDC20*-334 and *SPG20*-3570 did not meet our enrichment hypothesis (t-test: Mean ± SD; n = 3; * $p < 0.05$, ** $p < 0.01$ *** $p < 0.001$).

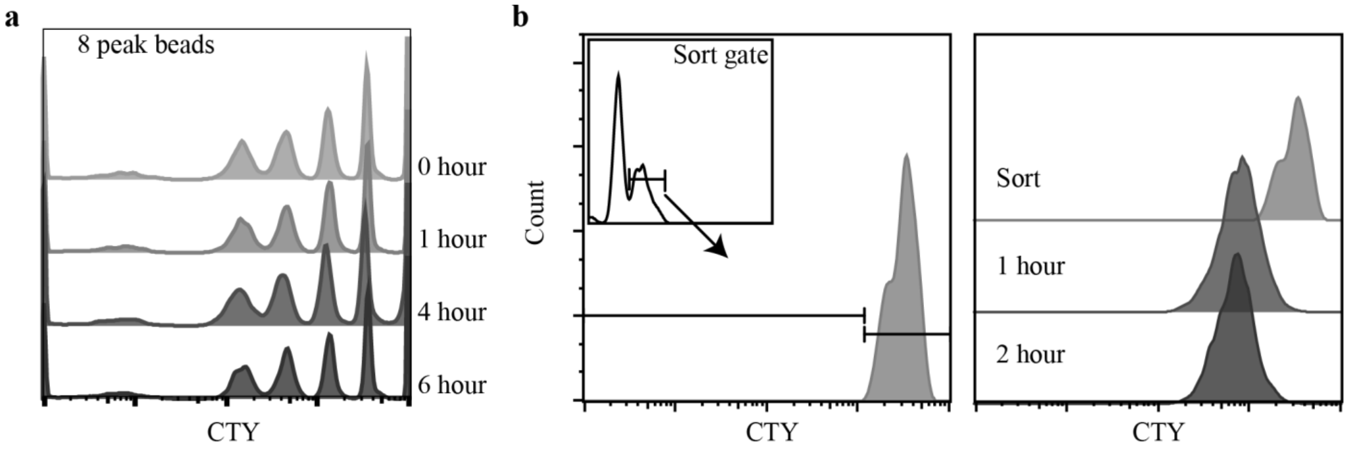


**Supplementary Figure 2. Flow cytometric analysis of intrinsic and extrinsic error associated with CTY proliferation assays. a)** To monitor machine (extrinsic) related error during CellTrace time course experiments we ran standardized 8 peak beads prior to each measurement to ensure that fluorescent peaks from the sorted parameter (CTY) were the consistent between each time point. Representative flow cytometric analyses of 8 peak bead CTY fluorescence showed limited signal shifts between 0 and 6 h. **b)** We observed an efflux of CellTrace Yellow (even after titration) in all experiments during the proliferation time course. We added an additional biological control in an effort to observe the same phenomenon. hiPSC-CM D8 cells were treated with Nocodazol for 48 hours and arrested at G2/M. Cells were dissociated to single cell suspensions and the G2/M peak events were sorted and replated for 1 h and 2 h. An observed decrease in CTY fluorescence at 1 h was evident in arrested cells and was attributed to a process other than cellular division. Note: Nocodazol was kept in suspension throughout sort and replating.

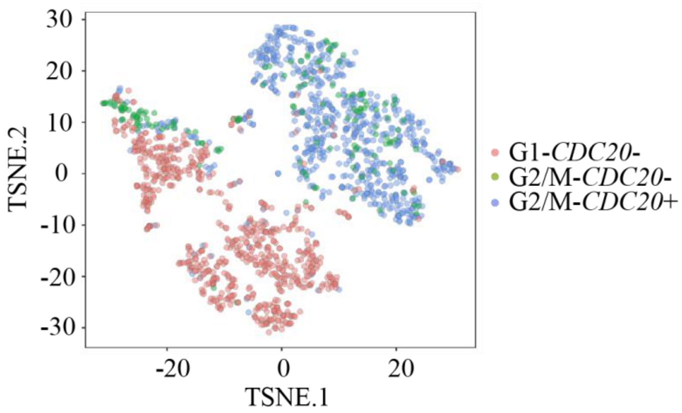


**Supplementary Figure 3. tSNE visualization of scRNA-seq data from each sorted population.**

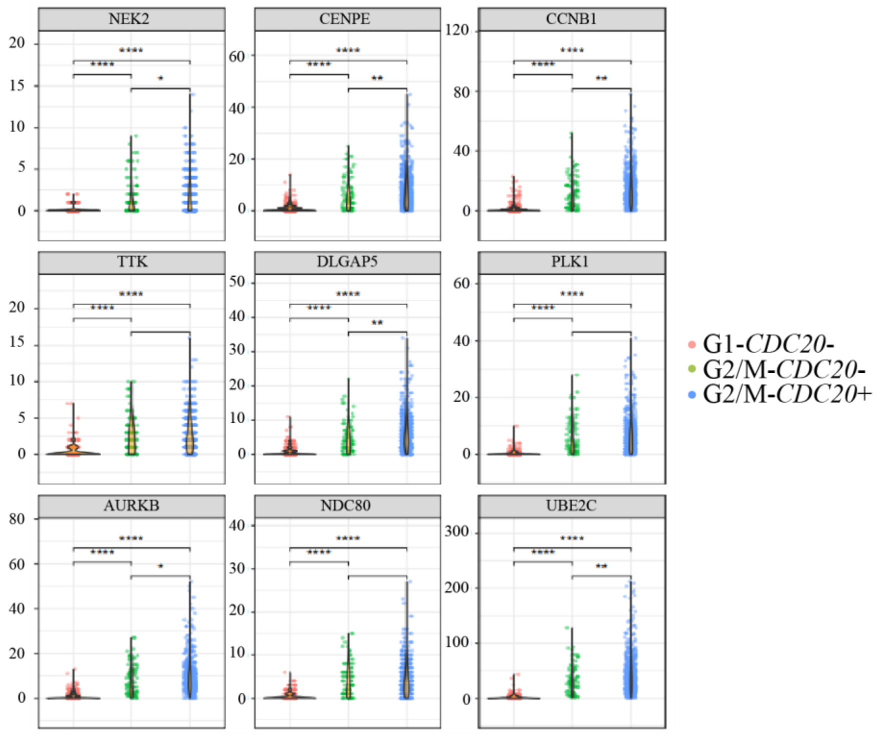


**Supplementary Figure 4. Genes expressed in G2/M-*CDC20*$^{+}$ cells that occupy the largest number of upregulated GO terms.** Violin plots generated using the pagoda2 package to identify genes that were differentially expressed between G1-*CDC20*$^{-}$ (orange), G2/M-*CDC20*$^{-}$ (green) and G2/M-*CDC20*$^{+}$ (blue) cell populations based on single cell expression levels. Data sets were filtered to include cells with > 500 transcripts and genes > 10 transcripts across all cells. Data were normalized to reduce expression level-dependent bias in variance. (t-test: Mean expression * $p < 0.05$ ** $p < 0.01$ **** $p < 0.0001$).

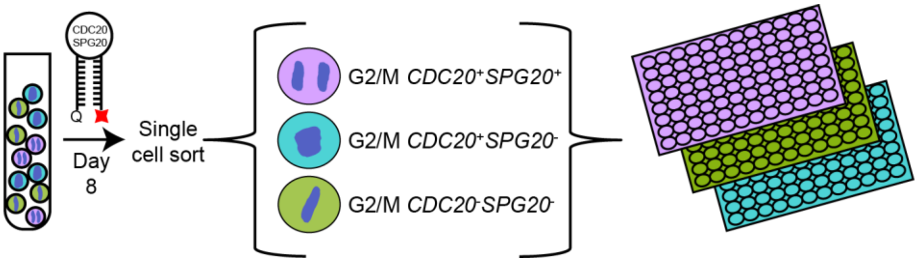


**Supplementary Figure 5. Schematic of the protocol used for delivery of *CDC20* and *SPG20* MBs, isolation, and quantification of nuclei morphology of single sorted events from D8 hiPSC-CMs in 96 well plates.**

**Supplementary Table 1. Design of molecular beacons**

| MB | 5' Probe | 3' Quencher | MB sequence (5'-3')* | Target sequence (5'-3') |
|---|---|---|---|---|
| *CDC20-642* | Quasar670 | BHQ2 | CCTGGGTAATAGTCATTTCGGATCCAGG | ATCCGAAATGACTATTAC |
| *SPG20-2264* | Cy3 | BHQ2 | CAGAGCTTTCTTCTTTGCCTCCCTCTG | GGAGGCAAAGAAGAAAG |
| Negative control MB | Cy3 or Quasar670 | BHQ2 | CGCGACGACAAGCGCACCGATATCGCG | |
| Positive control MB | Cy3 or Quasar670 | n/a | CGCGACGACAAGCGCACCGATATCGCG | |

*Underlined bases indicate stem sequences

**Supplementary Table 2. Sorted cell population names and corresponding flow cytometry parameters**

| **Instrument: BD Influx** | | | | | |
|---|---|---|---|---|---|
| **Laser Lines** | 640 | 488 | 405 | 640 | 561 |
| **Emission Filters** | 750LP | 530/40 | 460/50 | 670/30 | 585/29 |
| **Fluorochrome** | fvd780 | SIRPA FITC | DCV | *CDC20* Q670 | *SPG20* Cy3 |
| **G1-*CDC20*$^{-}$** | - | high | G1 | - | n/a |
| **G2/M-*CDC20*$^{-}$** | - | high | G2/M | - | n/a |
| **G2/M-*CDC20*$^{+}$** | - | high | G2/M | + | n/a |
| **G1-*CDC20*$^{-}$*SPG20*$^{-}$** | - | high | G1 | - | - |
| **G2/M-*CDC20*$^{-}$*SPG20*$^{-}$** | - | high | G2/M | - | - |
| **G2/M-*CDC20*$^{+}$*SPG20*$^{-}$** | - | high | G2/M | + | - |
| **G2/M-*CDC20*$^{+}$*SPG20*$^{+}$** | - | high | G2/M | + | + |

**Supplementary Table 3. Differentially expressed genes based on single cell expression levels**

| Genes | $log_2(FC)$ | $-log_{10}(FDR)$ |
|---|---|---|
| Upregulated | | |
| UBE2C | 0.71457 | 75.48435 |
| KPNA2 | 0.60798 | 74.71023 |
| TUBB4B | 0.59737 | 69.27450 |
| AURKB | 0.68055 | 67.45319 |
| TOP2A | 0.66683 | 63.19961 |
| PLK1 | 0.72972 | 59.21775 |
| CCNB1 | 0.64122 | 58.89435 |
| TPX2 | 0.65563 | 56.42611 |
| CDC20 | 0.65919 | 54.84472 |
| SGOL2 | 0.69749 | 53.44951 |
| CCNA2 | 0.60613 | 51.41900 |
| AURKA | 0.72380 | 51.16399 |
| FAM64A | 0.60956 | 51.09119 |
| DLGAP5 | 0.64388 | 51.09119 |
| CDCA3 | 0.72680 | 50.61288 |
| NUF2 | 0.59791 | 50.42378 |
| HMMR | 0.67589 | 49.65935 |
| GTSE1 | 0.63603 | 48.96168 |
| CENPA | 0.70209 | 48.88983 |
| UBE2S | 0.61338 | 48.23050 |
| DEPDC1 | 0.69705 | 47.93384 |
| ASPM | 0.65624 | 47.58927 |
| MKI67 | 0.61154 | 47.54783 |
| CDCA8 | 0.65172 | 47.40499 |
| CENPE | 0.61022 | 46.71888 |
| NDC80 | 0.66737 | 46.31268 |
| PSRC1 | 0.65719 | 44.70072 |
| KNSTRN | 0.61047 | 44.46471 |
| PRC1 | 0.61376 | 41.86038 |
| TTK | 0.62583 | 36.61727 |
| CDKN3 | 0.59621 | 36.36420 |
| PIF1 | 0.72434 | 34.99925 |
| KIF23 | 0.60510 | 33.42616 |
| FAM83D | 0.64634 | 33.11878 |
| CKAP2L | 0.69315 | 32.45788 |
| BUB1 | 0.61610 | 30.19732 |
| KIF2C | 0.60353 | 28.60805 |
| CDC25C | 0.67853 | 28.19648 |
| ARHGAP11A | 0.60119 | 28.09073 |
| RACGAP1 | 0.60821 | 27.52537 |

| | | |
|---|---|---|
| NEK2 | 0.72496 | 25.23105 |
| KIF14 | 0.63010 | 24.39957 |
| ECT2 | 0.58929 | 23.67704 |
| FAM72C | 0.74352 | 16.94356 |
| FAM72D | 0.68388 | 14.85591 |
| BORA | 0.63329 | 14.59604 |
| GAS2L3 | 0.75059 | 12.69158 |
| TNFAIP8L1 | 0.59789 | 3.35506 |
| APOLD1 | 0.67224 | 3.18546 |
| Down regulated | | |
| PAQR4 | -0.96873 | 3.01529 |
| MRPL53 | -0.63580 | 3.02091 |
| CDC6 | -0.78162 | 3.31775 |
| CHAF1B | -0.75979 | 3.97115 |
| WDR76 | -1.03357 | 5.25855 |
| SNHG19 | -0.72872 | 7.54460 |
| FAM111B | -1.13629 | 7.66483 |
| ATP5L2 | -0.59729 | 7.83900 |
| NME2 | -0.59789 | 8.25299 |
| CDCA7 | -0.85415 | 8.65230 |
| MCM2 | -0.62889 | 10.14641 |
| SNHG25 | -0.73303 | 11.02485 |
| TIPIN | -0.62007 | 15.79345 |
| UHRF1 | -1.10787 | 17.93307 |
| MCM4 | -0.92443 | 18.35493 |
| MCM6 | -1.29305 | 23.67704 |
| PCNA | -0.75998 | 24.74748 |
| MCM3 | -0.63614 | 24.95847 |
| MCM5 | -1.10653 | 25.26889 |
| UNG | -1.15725 | 27.86425 |
| NACA2 | -0.69511 | 34.35303 |
